# Combined multiple transcriptional repression mechanisms generate ultrasensitivity and oscillations

**DOI:** 10.1101/2022.01.19.476033

**Authors:** Eui Min Jeong, Yun Min Song, Jae Kyoung Kim

**Affiliations:** Department of Mathematical Sciences, Korea Advanced Institute of Science and Technology, Daejeon 34141, Republic of Korea; Biomedical Mathematics Group, Institute for Basic Science, Daejeon 34126, Republic of Korea

**Keywords:** Transcriptional repression, sequestration, displacement, ultrasensitivity, biological oscillators, circadian clock

## Abstract

Transcriptional repression can occur via various mechanisms, such as blocking, sequestration, and displacement. For instance, the repressors can hold the activators to prevent binding with DNA or can bind to the DNA-bound activators to block their transcriptional activity. Although the transcription can be completely suppressed with a single mechanism, multiple repression mechanisms are utilized together to inhibit transcriptional activators in many systems, such as circadian clocks and NF-κB oscillators. This raises the question of what advantages arise if seemingly redundant repression mechanisms are combined. Here, by deriving equations describing the multiple repression mechanisms, we find that their combination can synergistically generate a sharply ultrasensitive transcription response and thus strong oscillations. This rationalizes why the multiple repression mechanisms are used together in various biological oscillators. The critical role of such combined transcriptional repression for strong oscillations is further supported by our analysis of formerly identified mutations disrupting the transcriptional repression of the mammalian circadian clock. The hitherto unrecognized source of the ultrasensitivity, the combined transcriptional repressions, can lead to robust synthetic oscillators with a previously unachievable simple design.

## 1. Introduction

Transcription, the first step of gene expression, is regulated by activators and repressors, i.e., the bindings of the activators and repressors to a specific DNA sequence promote and down-regulate transcription, respectively [1, 2]. The repressors can also indirectly inhibit transcription by binding with the activators rather than with DNA (figure 1a) [3, 4]. That is, the repressors can bind to the DNA-bound activators to block their transcriptional activity (Blocking; figure 1a), hold the activators to prevent them from binding with DNA (Sequestration; figure 1a), and dissociate the activators from DNA by forming a complex (Displacement; figure 1a).

**Figure 1.**
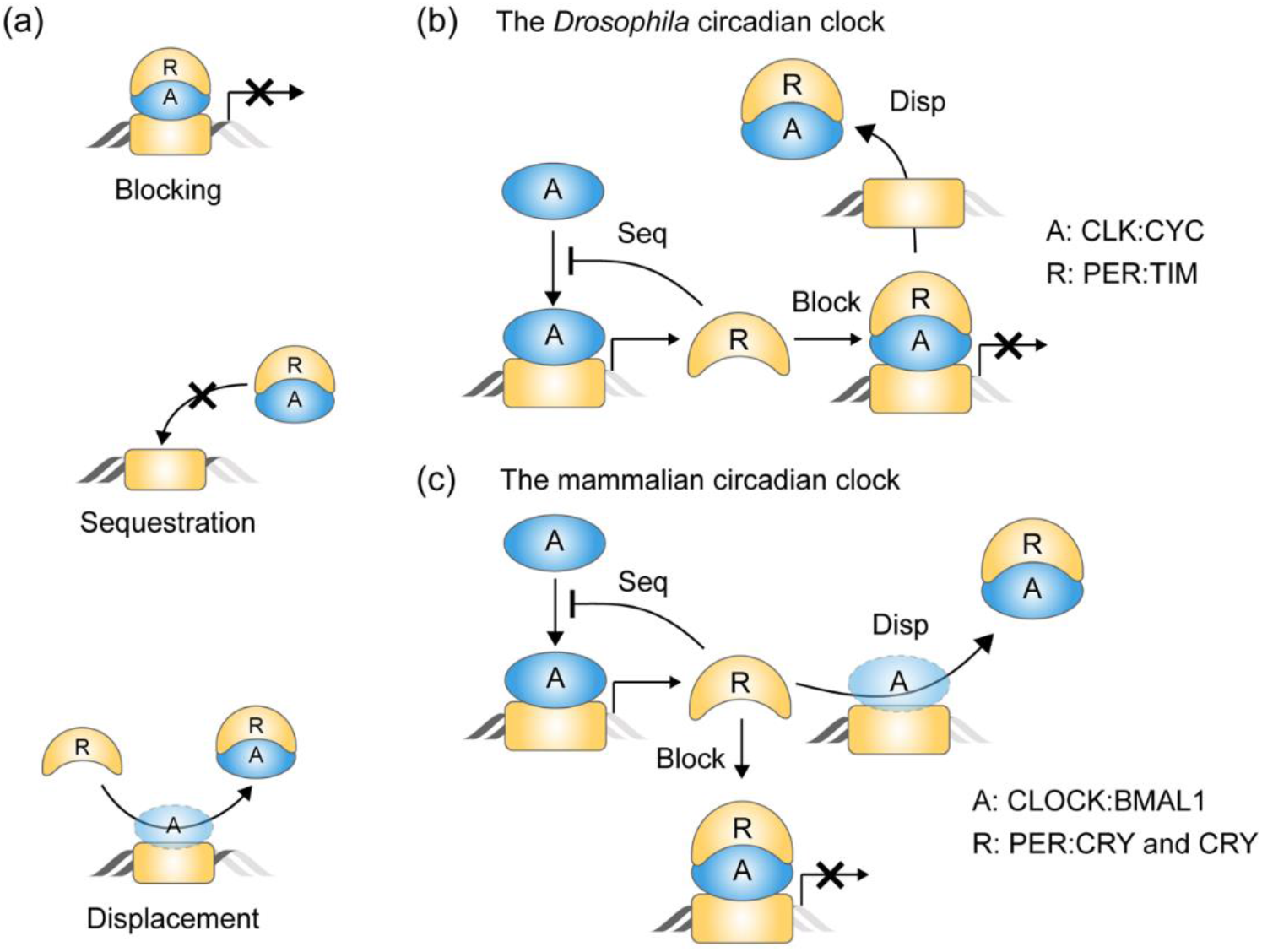
Multiple transcriptional repression mechanisms are used together in the transcriptional NFL of the circadian clock. **(a)** Repressors (*R*) can suppress the transcriptional activity of activators (*A*) with multiple mechanisms. For example, *R* binds to the DNA-bound *A* to block its transcriptional activity (Blocking), holds *A* to prevent binding to DNA (Sequestration), or dissociates *A* from DNA (Displacement). **(b)** Such multiple repression mechanisms are used together in the transcriptional NFL of the *Drosophila* circadian clock. The activator CLK:CYC (*A*) promotes the synthesis of the repressor PER:TIM (*R*). Then, *R* inhibits the transcriptional activity of *A* in various ways: *R* sequesters the free *A* from DNA, and blocks the transcriptional activity of the DNA-bound *A* and then displaces it from DNA. **(c)** Similarly, in the mammalian circadian clock, the repressors PER:CRY and CRY (*R*) inhibit their own transcriptional activator CLOCK:BMAL1 (*A*) via blocking, sequestration, and displacement.

Each repression mechanism appears to be able to suppress transcription solely. However, various repressors utilize combinations of multiple repression mechanisms [3]. For example, retinoblastoma (Rb) protein, a key regulator of mammalian cell cycle genes, represses transcription by blocking the activator and recruiting histone deacetylase, which alters the structure of chromatin [5–7]. Similarly, PHO80, a component of a yeast nutrient-responsive signaling pathway, represses transcription by blocking the activator and sequestering the activator in the cytoplasm [8–10]. This raises the question of the advantages of utilizing a combination of multiple repression mechanisms, which seems redundant.

In the transcriptional negative feedback loop (NFL) of various biological oscillators, repressors also inhibit their own transcriptions via combinations of the multiple repression mechanisms. For example, I*κ*B*α* inhibits its own transcriptional activator NF-*κ*B by sequestering it in the cytoplasm [11] as well as displacing it from DNA [12, 13], which induces the NF-κB oscillation under stress conditions. In the transcriptional NFL of the circadian clock, the transcription is also suppressed in multiple ways. Specifically, in the *Drosophila* circadian clock, the repressor (PER:TIM) sequesters its own transcriptional activator (CLK:CYC) from DNA (sequestration), blocks the transcriptional activity by binding to DNA-bound CLK:CYC (blocking), and then displaces it from DNA (displacement; figure 1b) [14]. Similarly, in the mammalian circadian clock, the repressor (PER:CRY and CRY) also inhibits its own transcriptional activator (BMAL1:CLOCK) by sequestration, blocking, and displacement (figure 1c) [15–17].

The transcriptional NFL can generate oscillations when the transcriptional activity shows an ultrasensitive response to changes in the concentration of repressors [18–21]. Such ultrasensitivity can be generated solely by sequestration when the activators and repressors tightly bind [22, 23]. In particular, the sequestration requires only tight binding, which seems to be physiologically more achievable than the conditions for the other ultrasensitivity-generating mechanisms based on cooperativity (e.g., cooperative oligomerization). Thus, sequestration has recently been adopted for mathematical models of circadian clocks [21, 24–30]. However, Heidebrecht et al. pointed out that the tightness of the binding between the activator and repressor required for the sequestration to generate sustained rhythms is beyond the physiologically plausible binding affinity [31].

Here, we find that combining multiple transcriptional repression mechanisms can synergistically generate ultrasensitivity by deriving their governing equations. Specifically, we find that the sole blocking-type repression can generate only low-sensitivity transcriptional activity. When sequestration is added, the ultrasensitivity can be generated only with strong blocking and stronger sequestration compared to the blocking. The required strong sequestration is challenging to achieve with a physiologically plausible binding affinity. Interestingly, this limitation to generate ultrasensitivity and strong oscillations can be overcome by adding displacement. To test whether the combination of the multiple repressions is critical for the mammalian circadian clock to generate strong rhythms, we investigated the previously identified mutations disrupting the transcriptional repressions [32–35]. Indeed, when any of the blocking, sequestration, or displacement was disrupted, the circadian rhythms of PER2-LUC became weaker in mice. Our work explains why the combination of seemingly redundant repression mechanisms is used in various systems requiring ultrasensitivity, such as the cell cycle and the circadian clock.

## 2. Results

### 2.1. The sole blocking-type repression generates a hyperbolic response in the transcriptional activity

To investigate how the transcription is regulated by the multiple repression mechanisms (figure 1a), we first constructed a model describing the single blocking-type repression (figure 2a; see Methods for details). In the model, the transcription is triggered when the free activator (*A*) binds to the free DNA (*E_F_*) with a dissociation constant of *K_a_* to form the activated DNA (*E_A_*). The transcription is inhibited when the repressor (*R*) binds to the DNA-bound *A* (*E_A_*) to form ternary complex (*E_R_*) with a dissociation constant of *K_b_* (i.e., the blocking-type repression). Therefore, the transcriptional activity is proportional to the probability that DNA is bound with only *A* and not *R*, i.e., *E_A_/E_T_*, where *E_T_* = *E_F_* + *E_A_* + *E_R_* is the conserved total concentration of DNA. In particular, when the transcription rate is normalized to one, the transcriptional activity and *E_A_/E_T_* become the same. Thus, for simplicity, we refer to *E_A_/E_T_* as the transcriptional activity throughout this study.

**Figure 2.**
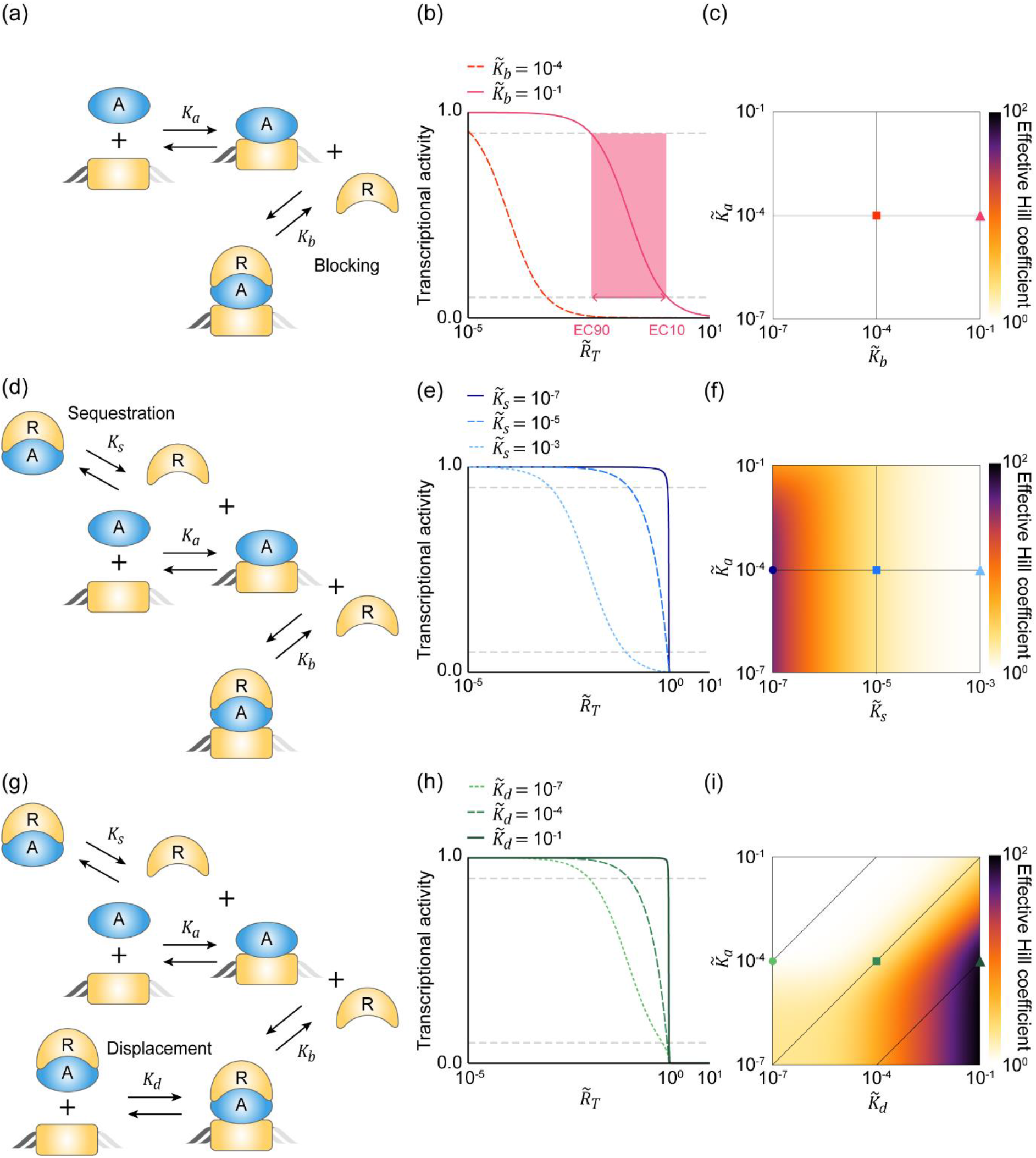
The combination of multiple repression mechanisms leads to ultrasensitive transcription response. **(a)** Diagram of the model describing the blocking-type repression. The binding of the activator (*A*) to DNA with a dissociation constant of *K_a_* leads to the transcription, and the binding of the repressor (*R*) to the DNA-bound *A* with a dissociation constant of *K_b_*, inhibits the transcription. **(b)** As the molar ratio between the total repressor and activator concentrations 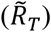 increases or their binding affinity increases (i.e., 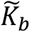 decreases), the transcriptional activity decreases. The sensitivity of the transcriptional activity is quantified using the effective Hill coefficient 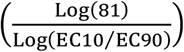, which increases as the width of the EC90 and EC10 box decreases (i.e., the red box). The grey dashed lines denote the 10% and 90% values of the maximal transcriptional activity, respectively. Here, 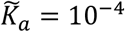. (**c**) The effective Hill coefficient is one regardless of the values of 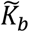 and 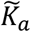, indicating that the sole blocking can generate only low sensitivity. The square and triangle marks represent the parameter values used for (b). **(d)** The sequestration-type repression is added to the blocking model in (a): *R* sequesters the free A with a dissociation constant of *K_s_* from DNA. **(e)** When the sequestration is weaker than the blocking (i.e., 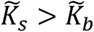; dotted line), the sensitivity of the transcriptional activity is similar to that regulated by only the blocking-type repression (b). On the other hand, when the sequestration is stronger than the blocking (i.e., 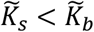; solid and dashed lines), a switch-like transition in the transcriptional activity occurs. Here, 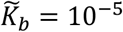 and 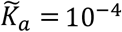. (**f**) The effective Hill coefficient increases as 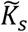 decreases. The circle, square, and triangle marks represent the parameter values used for (e). **(g)** The displacement-type repression is added to the model in (d): the *R_A_* complex dissociates from DNA with a dissociation constant of *K_d_*. **(h)** When 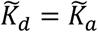 (dashed line), it satisfies the detailed balance condition (i.e., 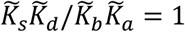) and thus the displacement has no effect on the transcriptional activity (cf. dashed line in (e)). When the effective displacement occurs (i.e., 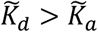; solid line), the sensitivity increases. Here, 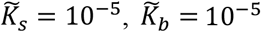 and 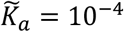. **(i)** When 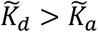, the effective Hill coefficients become extremely larger compared to those obtained with the sequestration and blocking (f). The circle, square, and triangle marks represent the parameter values used for (h).

The transcriptional activity (*E_A_/E_T_*) increases as *A* increases or *R* decreases. This relationship can be quantified by deriving the steady state of *E_A_/E_T_*. Because the steady state of *E_F_* depends on the single pair of binding and unbinding reactions with the dissociation constant of *K_a_*, its steady state equation is *AE_F_* = *K_a_E_A_*. Similarly, the steady state equation of *E_R_* is also simple as *RE_A_* = *K_b_E_R_*. Therefore, 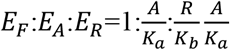 at the steady state, leading to the steady state of *E_A_/E_T_* as follows:

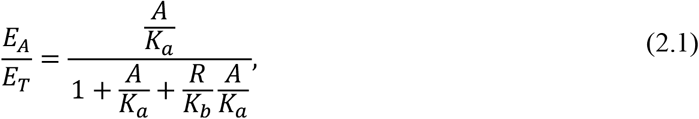

where *A* and *R* are the steady states of the free activator and repressor, respectively (see Methods for details). Because the steady states of *A* and *R* depend on the dissociation constants (i.e., *K_a_* and *K_b_*), it is challenging to analyze equation (2.1).

Equation (2.1) can be further simplified because the concentration of DNA is typically negligible compared to the concentration of activator and repressor proteins (see Methods for details). Specifically, *E_A_* and *E_R_* can be neglected in the conserved total concentration of the activator (*A_T_* = *A* + *E_A_* + *E_R_*) and the repressor (*R_T_* = *R* + *E_R_*), and thus *A* ≈ *A_T_* and *R* ≈ *R_T_*. This allows us to get the simplified approximation for equation (2.1) as follows:

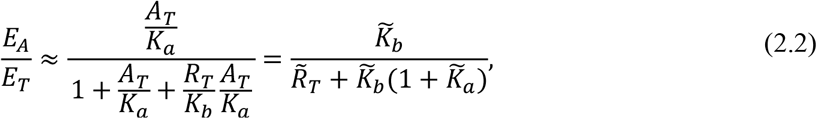

where 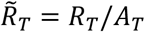 is the molar ratio between *R_T_* and *A_T_*, and 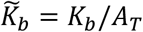 and 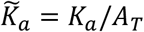 are the dissociation constants normalized by the concentration of the total activator. Equation (2.2) indicates that the transcriptional activity shows a hyperbolic response with respect to the molar ratio 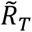 (figure 2b). Specifically, when 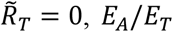 has the maximum value 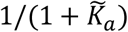, which becomes closer to one as *A* binds to DNA more tightly (i.e., 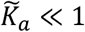). When 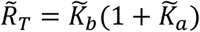, *E_A_/E_T_* is reduced to its half-maximal value. Thus, as the binding between the DNA-bound *A* and *R* becomes tighter (i.e., 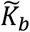 decreases), the transcriptional activity achieves its half-maximal value at the lower 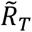 (figure 2b). The sensitivity of the transcriptional activity with respect to 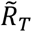 can be quantified using the effective Hill coefficient 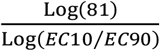, which is equivalent to the Hill exponent for a Hill curve [36]. The effective Hill coefficient of the transcriptional activity is one regardless of the 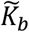 and 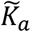 values (figure 2c), as expected from the Michaelis-Menten-type equation (equation (2.2)). Taken together, with the sole blocking repression, the transcriptional activity cannot sensitively respond to 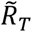.

### 2.2. The combination of the sequestration- and blocking-type repressions can generate the ultrasensitivity

We wondered whether the sensitivity of the transcriptional activity can be increased by incorporating an additional repression mechanism. To investigate this, we added the sequestration-type repression to the blocking model: *R* binds with the free *A* to form complex *R_A_* with a dissociation constant of *K_s_*, and thus sequesters *A* from DNA (figure 2d; see Methods for details). Due to the complex *R_A_*, the conservations are switched to *A_T_* = *A* + *R_A_* + *E_A_* + *E_R_* and *R_T_* = *R* + *R_A_* + *E_R_*. When the binding between *A* and *R* is weak (i.e., 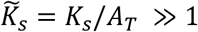) and thus *R_A_* is negligible, the steady states of *A* and *R* can be approximated with simple *A_T_* and *R_T_*. On the other hand, when the binding is not weak, *R_A_* is not negligible and thus the approximations for the steady states of *A* and *R* become a bit complex (see Methods for details):

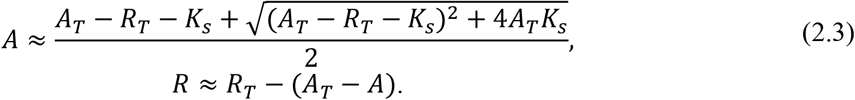

When the binding between *A* and *R* is extremely tight 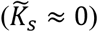, *A* and *R* can be approximated by the simple functions max(*A_T_* – *R_T_*, 0) and max(*R_T_* – *A_T_*, 0), respectively [21, 28, 30]. By substituting equation (2.3) for equation (2.1), the approximated *E_A_/E_T_* can be derived:

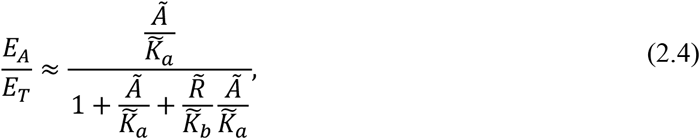

where 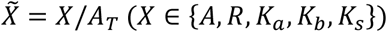. Because 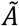 and 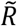 are determined by the molar ratio 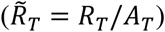, *E_A_/E_T_* is still the function of 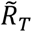 like in equation (2.2).

The transcriptional activity described by equation (2.4) shows more sensitive responses with respect to 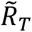 compared to the blocking model as the sequestration becomes stronger (i.e., 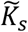 decreases; figure 2e). Specifically, when the sequestration is weaker than the blocking (i.e., 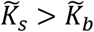), the transcriptional regulation is mainly governed by the blocking, and thus the transcriptional activity shows a hyperbolic response (figure 2e, dotted line) similar to the sole blocking-type repression (figure 2b). On the other hand, when the sequestration is stronger than the blocking (i.e., 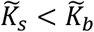; figure 2e, solid line), *R* is more likely to bind with the free *A* rather than the DNA-bound *A*. Thus, when there are more activators than repressors (i.e., 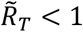), the majority of *R* is bound to the free *A*, not the DNA-bound *A*, and thus the high level of transcriptional activity is maintained. As 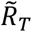 is greater than one and thus the free *R*, not sequestered by the free *A*, is available, *R* can block the DNA-bound *A*, leading to the rapid drop in the transcriptional activity (figure 2e, solid line). This switch-like transition in the transcriptional activity generates the ultrasensitivity (figure 2e). Consistently, the effective Hill coefficient increases as the sequestration becomes stronger (i.e., 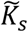 decreases; figure 2f).

The ultrasensitivity can be generated when the blocking and sequestration act synergistically (figure S1). That is, when the blocking is stronger than the sequestration (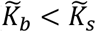; figure S1a-c), the ultrasensitivity cannot be generated, similar to the sole blocking model (figure 2c). When the blocking is too weak 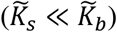, and thus the transcriptional regulation is mainly governed by the sequestration, the DNA-bound activator cannot be inhibited effectively via blocking. As a result, ultrasensitivity cannot be generated when the activator binds to DNA more tightly than the repressor (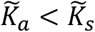; figures S1d and e). Taken together, to generate ultrasensitivity, the appropriate level of blocking and the stronger sequestration compared to the blocking are needed. This requires a mechanism for the repressor to have different binding affinities with the free activator and the DNA-bound activator. Furthermore, due to the requirement of stronger sequestration compared to the blocking, the condition is challenging to achieve with physiologically plausible binding affinities. Specifically, the concentration of transcriptional factors (*A_T_*) is 2 × 10^−9^~10^−7^ *M* as their number is 10^4^~10^5^ (i.e., 2 × 10^−20^ ~10^−19^ *mol*) and the typical mammalian cell volume is 10^−11^~10^−12^ *L* [31, 37, 38]. Thus, even the extremely high affinity protein whose dissociation constant is picomolar (i.e., *K_s_* ≈10^−12^ *M*) has the 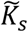 with the range of 0.5 × 10^−5^~10^−3^. With these physiologically plausible values of 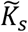, the range of 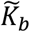 where the ultrasensitivity can be generated is narrow (figure S1).

### 2.3. The combination of the displacement-, sequestration- and blocking-type repressions can readily generate the ultrasensitivity under physiologically plausible conditions

To investigate whether the requirement of the strong sequestration can be relaxed by adding the displacement-type of repression, we expanded the model where the complex *R_A_* can dissociate from DNA with a dissociation constant of *K_d_* (figure 2g; see Methods for details). Due to the displacement, *E_F_* is affected by two different reversible bindings between *R_A_* and *E_F_* as well as between *A* and *E_F_* unlike in the previous models. Thus, the steady state equation of *E_F_* is switched to *AE_F_* + *R_A_E_F_* = *K_a_E_A_* + *K_d_E_R_* from *AE_F_* = *K_a_E_A_* (see Methods for details). Similarly, the steady state equation of *E_R_* is also switched to *K_b_E_R_* + *K_d_E_R_* = *RE_A_* + *R_A_E_F_* from *K_b_E_R_* = *RE_A_*. By solving these coupled equations, we can get the ratio of the steady states of *E_F_*, *E_A_*, and *E_R_*, i.e., 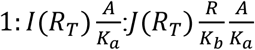, where 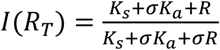 and 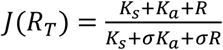, and 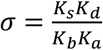. Note that when *σ* = 1, which is known as the detailed balance condition [39], *I*(*R_T_*) = *J*(*R_T_*) = 1 and thus the ratio becomes the same as the previous simple one. This is because under the detailed balance condition, all reversible bindings reach equilibrium, and thus the steady state equations of the species affected by multiple reversible reactions (e.g., *AE_F_* + *R_A_E_F_* = *K_a_E_A_* + *K_d_E_R_*) can be partitioned into the equilibrium relations for each reversible reaction (i.e., *AE_F_* = *K_a_E_A_* and *R_A_E_F_* = *K_d_E_R_*) [39]. Therefore, under the detailed balance condition, the transcriptional repression by the three types of repressions becomes equivalent to the repression by the blocking- and sequestration-types.

When *σ* ≠ 1, the ratio of the steady states of *E_F_*, *E_A_*, and *E_R_* are changed and thus we get 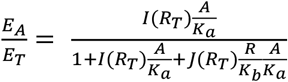, different from equation (2.1). After substituting equation (2.3) into *A* and *R* and normalizing the variables and parameters with *A_T_*, we can derive the approximation for *E_A_/E_T_* in terms of the molar ratio 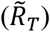:

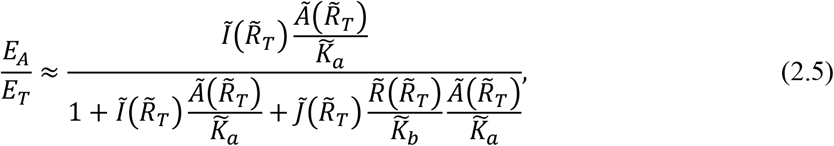

where 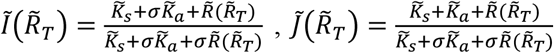 and 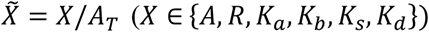. To investigate whether the displacement enhances the sensitivity of the transcriptional activity, we first consider the case where *R* binds to the free *A* and the DNA-bound *A* with the same affinity (i.e., 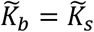), so the combination of the sequestration- and blocking-type repressions fails to generate the ultrasensitivity (figures 2f and S1). In this case, if *R_A_* and *A* have the same binding affinity with DNA (i.e., 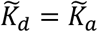), *σ* = 1 and thus the displacement-type repression does not make any difference compared to the combination of the sequestration- and blocking-type repressions (figure 2e and h, dashed lines). On the other hand, if effective displacement occurs (i.e., *R_A_* more easily dissociates from DNA compared to *A*, 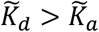), *E_R_* decreases and *E_A_* increases. As a result, the higher level of transcriptional activity is maintained until 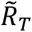 becomes closer to one, yielding greater sensitivity (figure 2h, solid line). Consistently, the effective Hill coefficient becomes larger as 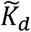 becomes greater than 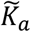 (figure 2i). Furthermore, even if 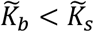 (i.e., the sequestration is weaker than the blocking), the ultrasensitivity can be generated when the effective displacement occurs (figure S2), unlike with the combination of blocking and sequestration (figure 2f and S1). Taken together, effective displacement can eliminate the requirement for the combination of the sequestration- and blocking-type repressions to generate the ultrasensitivity.

When there is no energy expenditure, the dissociation constants have to satisfy the detailed balance condition 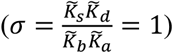 [40]. In this case, effective displacement can occur 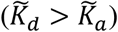 under limited conditions 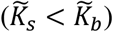, which is challenging to achieve physiologically. On the other hand, when energy is expended to break the detailed balance condition (*σ* > 1), effective displacement can occur without the limitation. Such energy expenditure can happen mechanistically by adenosine triphosphate hydrolysis [41].

Interestingly, when there is no energy expenditure to break the detailed balance condition, the equilibrium relations for each reversible reaction (i.e., *AE_F_* = *K_a_E_A_, R_A_E_F_* = *K_d_E_R_*) hold at the steady state [39]. Thus, the transcription regulated by all three repressions becomes the same as that regulated by any two of the repressions (see Supplementary Materials for details). This allows us to easily identify the condition for ultrasensitivity generated with any two repression mechanisms by substituting the detailed balance condition 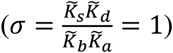 to the condition for the ultrasensitivity generated with the three repression mechanisms (table S1). This reveals that the requirement of strong sequestration of the blocking and sequestration model, which was challenging physiologically, is switched to the effective displacement of the blocking and displacement model. Importantly, with energy expenditure, the combination of all three repressions can generate ultrasensitivity over a wider range of condition compared to the combination of any two repressions (table S1, figure 2i, S2, and S3).

### 2.4. The transcriptional NFL with multiple repression mechanisms can generate strong rhythms

The ultrasensitivity is critical for the transcriptional NFL to generate sustained and strong oscillations [18–21]. Thus, when the transcriptional repression is regulated by the combination of the multiple repression mechanisms, the strong oscillations can be generated. To investigate this, we constructed a simple transcriptional NFL model (figure 3a), where the free activator (*A*) binds to the free DNA, and then promotes the transcription of the repressor mRNA (*M*). *M* is translated to the repressor protein in the cytoplasm (*R_c_*). After translocation to the nucleus, the repressor protein (*R*) inhibits its own transcriptional activator (*A*) with the previously described repression mechanisms (figure 2a, d, and g). Thus, the transcription of *M* depends on the transcriptional activity *E_A_/E_T_*. We assumed that *E_A_/E_T_* rapidly reaches its quasi-steady-state (QSS) because the reversible bindings regulating the transcriptional activity typically occur much faster than the other processes of the transcriptional NFL (i.e., transcription, translation, translocation, and degradation). Using the quasi-steady-state approximation (QSSA) and the non-dimensionalization, we can obtain the simple NFL model (see Supplementary Materials for details),

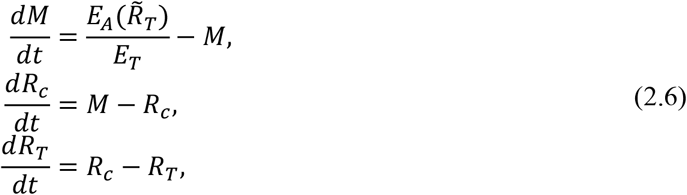

where 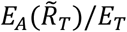 is the QSSA for the transcriptional activity. Depending on the repression mechanism, we can use the steady state equations for *E_A_/E_T_* derived in the previous sections (i.e., equations (2.2), (2.4), and (2.5)). Note that these QSSAs are known as the “total” QSSAs as they are determined by the molar ratio between the “total” concentrations of the repressor and activator, 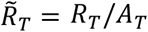, which is not affected by the fast reversible bindings. Thus, the QSSAs are accurate as long as the reversible bindings are fast [42]. In this way, the NFL model (equation (2.6)) can accurately capture the dynamics of the interactions between *A* and *R* even when their levels are comparable [42].

**Figure 3.**
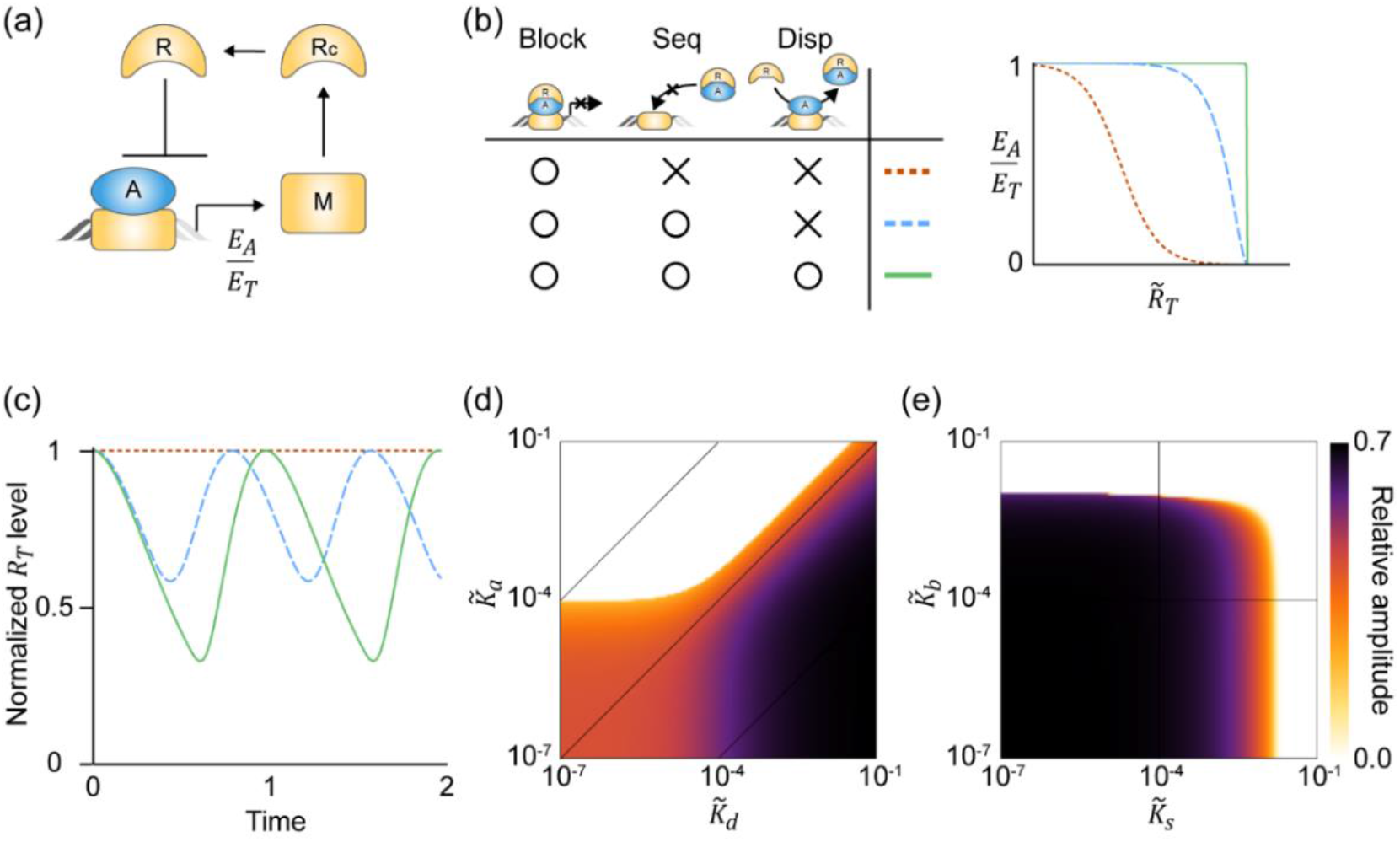
The transcriptional NFL with multiple repression mechanisms can generate strong oscillations. **(a)** In the transcriptional NFL model, the transcription rate of mRNA (*M*) is determined by the transcriptional activity (*E_A_/E_T_*). **(b)** *E_A_/E_T_* changes more sensitively in response to 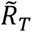 when more repression mechanisms are used, as shown in figure 2. **(c)** The NFL model with the sole blocking cannot generate rhythms (red dotted line), while the model with the multiple repressions can generate rhythms (blue dashed and green solid lines). In particular, the combination of the blocking, sequestration, and displacement can generate the strongest rhythms. Here, *A_T_* = 0.05, 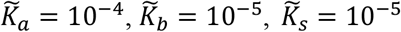, and 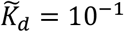 are used, and all trajectories are normalized by their own maximum value to compare their relative amplitudes. **(d)** As the displacement becomes ineffective (i.e., 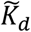 becomes smaller than 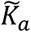), the relative amplitude decreases. **(e)** Similarly, as the blocking or sequestration becomes weaker (i.e., 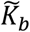 or 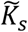 increases), the relative amplitude decreases.

As more repression mechanisms are added, 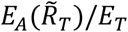 more sensitively changes in response to the variation of 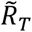 (figure 3b), which is critical for amplitude amplification. Thus, stronger rhythms, which have a high relative amplitude (i.e., the amplitude normalized by the peak value of the rhythm), are generated (figure 3c). Specifically, while the NFL with the sole blocking repression cannot generate rhythms (figure 3c, red dotted line), the NFL with the combination of the blocking, sequestration, and displacement can generate the strongest rhythms (figure 3c, green solid line). Such strong rhythms become weaker as the displacement becomes ineffective (i.e., 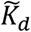 becomes smaller than 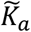; figure 3d), or the blocking or the sequestration become weaker (i.e., 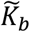 or 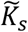 increases; figure 3e).

### 2.5. In the mammalian circadian clock, the disruption of synergistic multiple repressions weakens rhythms

In the transcriptional NFL of the mammalian circadian clock, the transcriptional repression occurs via the combination of blocking, sequestration, and displacement (figure 4a). Specifically, CLOCK:BMAL1 binding to E-box regulatory elements in the Period (Per1 and Per2) and Cryptochrome (Cry1 and Cry2) genes activates their transcription at around Circadian Time (CT) 4-8. After CRY and PER are translated in the cytoplasm, they form the complex with the kinase CK1δ and enter the nucleus. The complex dissociates CLOCK:BMAL1 from the E-box and sequesters CLOCK:BMAL1 to prevent binding to the E-box at around CT12-22 (displacement- and sequestration-type repression). At around CT0-4, CRY binds to the CLOCK:BMAL1:E-box complex to block the transcriptional activity (blocking-type repression) [15–17, 43].

**Figure 4.**
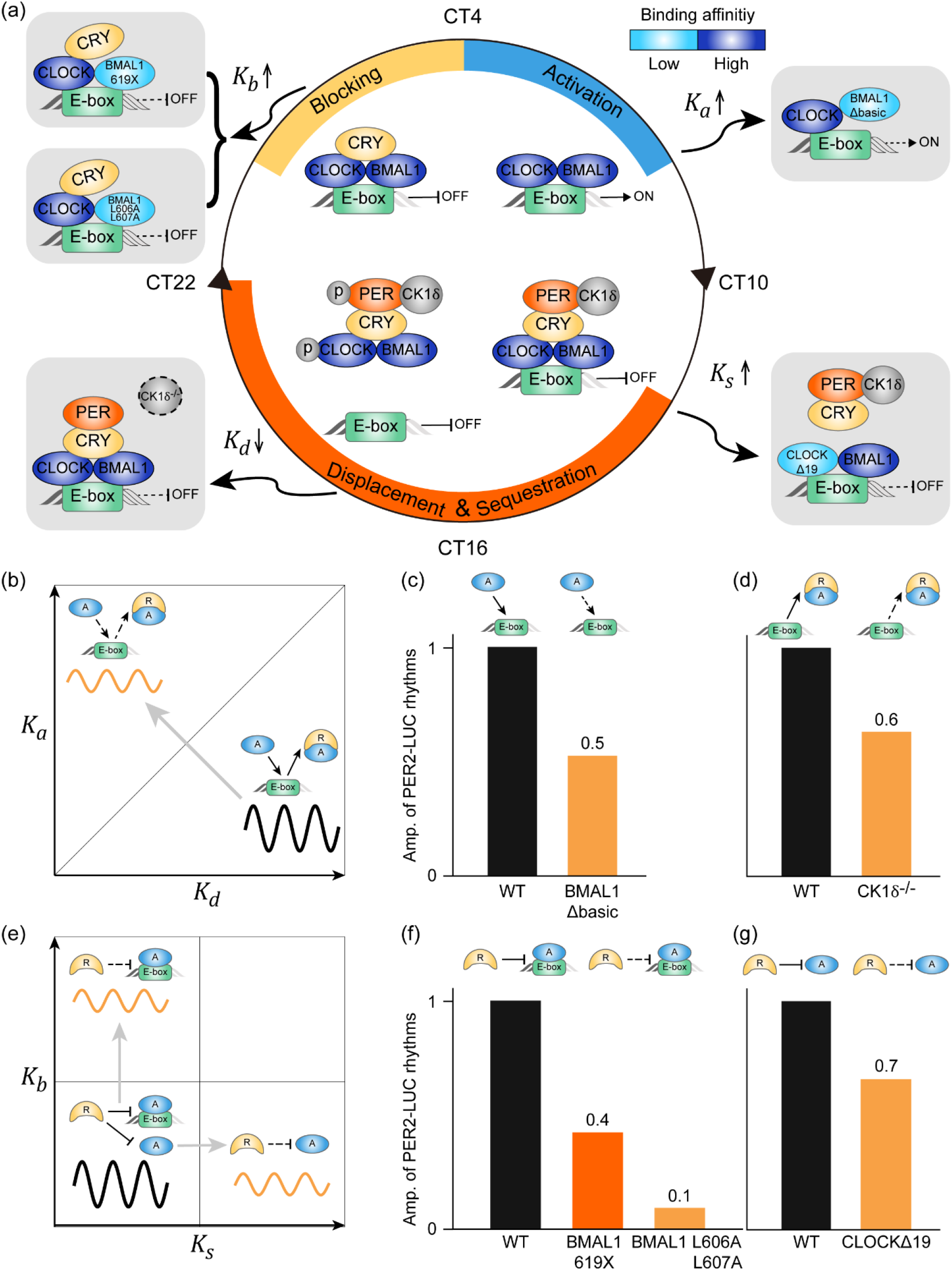
In the mammalian circadian clock, the disruption of synergistic multiple repressions weakens rhythms. **(a)** In the mammalian circadian clock, the transcriptional activity of CLOCK:BMAL1 is regulated by blocking-, sequestration-, and displacement-type repressions. Several mutations disrupting the combination of multiple repressions have been identified. BMAL1 transactivation domain mutations such as 619X and L606A L607A decrease the binding affinity between BMAL1 and CRY1 (i.e., *K_b_* increases), weakening the blocking. CLOCKΔ19 has impaired binding with PER (i.e., *K_s_* increase), disrupting the sequestration. A CK1δ^-/-^ mutation prevents the CK1δ-induced phosphorylation of CLOCK:BMAL1, which is essential for the effective displacement (i.e., *K_d_* decreases). Furthermore, BMAL1Δbasic has impaired binding with the E-box (i.e., *K_a_* increases), decreasing *K_d_/K_a_* and thus disrupting the effective displacement. **(b)** Schematic diagram showing the alteration of amplitudes by change in dissociation constants *K_a_* and *K_d_* based on the predictions in figure 3d. In the mammalian circadian clock, CLOCK:BMAL1 shows higher binding affinity with the E-box compared to the PER:CRY:CLOCK:BMAL1 complex (i.e., *K_d_/K_a_* > 1; below the gray line) [15], which is critical for strong rhythm generation according to our model prediction. **(c-d)** When *K_a_* was increased by the BMAL1Δbasic mutation (c) and *K_d_* was decreased by the CK1δ^-/-^ mutation (d), the amplitude of PER2-LUC rhythms was reduced to 0.5 and 0.6 compared to the WT mice, respectively. Adapted from [35] and [32]. **(e)** Schematic diagram showing the alteration of amplitudes by change in dissociation constants *K_s_* and *K_b_* based on the predictions in figure 3e. In the mammalian circadian clock, PER:CRY binds with CLOCK:BMAL1 tightly (i.e., small *K_s_*) and CRY binds with CLOCK:BMAL1:E-box tightly (i.e., small *K_b_*), which is crucial for strong rhythm generation according to our model prediction. **(f)** Indeed, as the dissociation constant between CRY and CLOCK:BMAL1:E-box (*K_b_*) was increased by the BMAL1 619X mutation, the amplitude of PER2-LUC rhythms from fibroblasts in mutant mice was reduced to 0.4 compared to that in WT. The amplitude was further reduced when the *K_b_* was further increased by the BMAL1 L606A L607A mutation. Adapted from [34]. **(g)** When the binding affinity between PER and CLOCK was decreased (i.e., *K_s_* increased) by the CLOCKΔ19 mutation, the amplitude of PER2-LUC rhythms in the SCN of mutant mice was reduced to 0.7 compared to that in WT mice. Adapted from [33]. For each mutation, all adapted PER2-LUC rhythms of WT and mutant mice were measured under the same condition. However, amplitudes among different mutations cannot be compared due to different experimental conditions.

In the mammalian circadian clock, because the PER:CRY complex recruits CK1δ, inducing dissociation of CLOCK:BMAL1 from the E-box [15], the binding affinity of CLOCK:BMAL with the E-box is higher compared to its complex with PER:CRY (i.e., *K_d_/K_a_ >* 1). This effective displacement is critical for strong rhythm generation (figure 4b, black solid line) according to our model prediction (figure 3d). Then we can expect that, as either *K_a_* increases or *K_d_* decreases, *K_d_/K_a_* decreases, which deactivates the displacement-type repression, the circadian rhythms become weaker (figure 4b; orange solid line). Indeed, when *K_a_* was increased by a BMAL1 mutant lacking the basic region (BMAL1Δbasic), which is critical for the binding of BMAL1 to the E-box element (figure 4a, top right), the amplitude of PER2-LUC rhythms from the fibroblasts of mutant mice was reduced compared to that from wild-type (WT) mice (figure 4c) [35]. Furthermore, when *K_d_* was decreased by a CK1δ^-/-^ mutant lacking the CK1δ-induced dissociation of CLOCK:BMAL1 from the E-box (figure 4a, bottom left), the amplitude of PER2-LUC rhythms in the suprachiasmatic nucleus (SCN) of mutant mice was also reduced compared to that in WT mice (figure 4d) [32]. Note that the amplitude reduction by the CK1δ^-/-^ mutant could be due to other factors because CK1δ also regulates the stability and nucleus entry of PER [44].

The blocking- and sequestration-type repressions also effectively occur in the mammalian circadian clock. That is, PER:CRY binds with CLOCK:BMAL1 tightly (i.e., small *K_s_*), and CRY binds with CLOCK:BMAL1:E-box tightly (i.e., small *K_b_*) [45]. Such tight bindings are important for strong rhythm generation (figure 4e, black solid line) according to our model prediction (figure 3e). Thus, as either *K_b_* or *K_s_* increases, weakening the blocking- or the sequestration-type repression, the rhythms are expected to become weaker (figure 4e, orange solid lines). Indeed, when *K_b_* was increased due to the BMAL1 619X mutation reducing the binding affinity between BMAL1 and CRY1 (figure 4a, top left), the amplitude of PER2-LUC rhythms from the fibroblasts of mutant mice was reduced to 0.4 compared to WT (figure 4f) [34]. When *K_b_* was further increased by a BMAL1 L606A L607A mutation, the amplitude was further reduced (figure 4f) [34]. Moreover, when *K_s_* was increased by the CLOCK mutant lacking the exon 19 region (CLOCKΔ19), which is required for the binding of PER (figure 4a, bottom right) [46], the amplitude of PER2-LUC rhythms in the SCN of mutant mice was reduced compared to that in WT mice (figure 4g) [33]. Note that such reduction of the amplitude by CLOCKΔ19 could be due to other factors such as the low transcriptional activity of CLOCKΔ19 [47] and the impaired binding with the E-box [48].

## 3. Discussion

Transcriptional repression plays a central role in precisely regulating gene expression [2]. Various mechanisms for the repression have been identified [2–4]. In particular, the transcriptional activators can be inhibited in various ways by repressors such as blocking, sequestration, and displacement (figure 1a). Interestingly, these repression mechanisms are used together to inhibit a transcriptional activator in many biological systems [3]. In this study, we found that multiple repression mechanisms can synergistically generate a sharply ultrasensitive transcriptional response (figure 2) and thus strong rhythms in the transcriptional NFL (figure 3). Consistently, the mutations disrupting any of the blocking, sequestration, or displacement in the transcriptional NFL of the mammalian circadian clock weaken the circadian rhythms (figure 4). Our work identifies a benefit of utilizing multiple repression mechanisms together, the emergence of ultrasensitive responses, which are critical for cellular regulation such as epigenetic switches, the cell cycle, and circadian clocks [22].

Recently, detailed transcriptional repression mechanisms underlying various biological systems have been identified. For instance, while MDM2 was known to inhibit p53 by promoting its degradation [49], recent studies have suggested that MDM2 can also inhibit p53 through displacement and blocking [50, 51]. In the Rb-E2F bistable switch, the suppressor Rb protein and the E2F family of transcription factors inhibit mutually with multiple repression mechanisms such as blocking and chromatin structure modification, which are critical to generate ultrasensitivity and thus the bistable switch of cell cycle [6, 7, 52]. However, such repression mechanisms have not yet been incorporated into the mathematical models [53–56]. Similarly, the recent discoveries of multiple repression mechanisms underlying biological oscillators such as the circadian clock [14–17] and the NF-κB oscillator [12, 13] have not been fully incorporated even in recent mathematical models [24, 57–62]. In particular, the majority of the mathematical models for various systems have used the simple Michaelis-Menten or Hill-type functions to describe the transcriptional repression regardless of its underlying repression mechanisms, which can distort the dynamics of the system [21, 42]. Our work highlights the importance of careful modeling of the transcriptional repression depending on blocking, sequestration, or displacement to accurately capture the underlying dynamics.

Interestingly, to fully utilize the three repression mechanisms, energy expenditure is required. Without the energy expenditure, the detailed balance condition needs to be satisfied 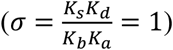. Under this restriction, the transcriptions regulated by the three repression mechanisms and any two of these become equivalent (see Supplementary Materials for details). As a result, the ultrasensitivity is generated under a limited condition compared to when the detailed balance condition is broken via dissipation of energy (*σ* > 1) (table 1, figures 2i, S1, and S2). Similarly, the limitation for generating the sensitivity of transcription under the detailed balance condition was also identified when DNA is directly regulated by its transcriptional factors [40]. Specifically, Estrada et al. found that when the energy expenditure breaks the detailed balance condition, the cooperative bindings of the transcriptional factors to multiple binding sites of DNA are more likely to generate ultrasensitivity.

The advantages of utilizing multiple repression mechanisms for biological oscillators have just begun to be investigated. For instance, in the NF-*κ*B oscillator, I*κ*B*α* inhibits its own transcriptional activator NF-*κ*B via sequestration and displacement. Z. Wang et al. found that the displacement can enhance NF-*κ*B oscillation by dissociating the NF-*κ*B from decoy sites and promoting its nuclear export (i.e., facilitating the sequestration), and compensating for the heterogeneous binding affinity of NF-*κ*B to the promoter of I*κ*B*α* [63]. Furthermore, a recent study of the transcriptional NFL of the mammalian circadian clock found that the displacement of the transcriptional activator (BMAL1:CLOCK) by its repressor (PER:CRY) can facilitate the mobility of the BMAL1:CLOCK to its various target sites, pointing out the hidden role of PER:CRY [64]. While PER:CRY dissociates and sequesters CLOCK:BAML1 from E-box (i.e., sequestration- and displacement-type), CRY blocks the transcriptional activity of CLOCK:BMAL1 (i.e., blocking type) [15–17]. Because Cry1 displays a delayed expression phase compared to Per, the blocking repression occurs at the late phase, which turns out to be critical for rhythm generation [65–67]. It would be interesting in future work to extend the model to include multiple repressors (e.g., PER and CRY) to investigate their distinct roles.

While we focused on transcriptional repression mechanisms, other mechanisms leading to the ultrasensitivity [68], and thus generating rhythms have been identified. For instance, phosphorylation of the repressor [24, 69, 70] and saturated degradation of the repressor [25, 31, 71] can be additional sources of ultrasensitivity for strong rhythms. Furthermore, an additional transcriptional positive feedback loop has been known to enhance the robustness of rhythms [18, 21, 70, 72] in the presence of Hill-type transcriptional repression, which can be induced by phosphorylation-based transcriptional repression [73, 74]. On the other hand, when the transcription is regulated by sequestration-type repression, an additional negative feedback loop rather than the positive feedback loop can enhance the robustness of rhythms [21, 28, 31]. It would be important in future work to investigate the role of additional feedback loops depending on the transcriptional repression mechanisms identified in this study.

A transcriptional NFL, where a single repressor inhibits its own transcription by binding to its own promoter, is the simplest design of the synthetic genetic oscillator [75, 76]. To generate the ultrasensitivity with this simple design, Stricker et al. used a repressor that forms a tetramer to bind with its own promoter [77]. Nonetheless, the degree of the ultrasensitivity was not enough for the synthetic oscillator to generate strong oscillations with high amplitude. Thus, more complex designs of synthetic oscillators have been constructed [75, 76]: the modified repressilators [78, 79], the combination of the negative and positive feedback loops [77, 80], and the coupling of synthetic microbial consortiums [81–84]. Our study proposes that a strong synthetic oscillator with a simple design (i.e., a single NFL) could be constructed by modifying the previously used repression mechanisms. That is, by using the combining blocking-, sequestration- and displacement-type repressions, although this might be challenging to implement, ultrasensitivity to achieve strong rhythms could be obtained, providing a new strategy for the design of synthetic oscillators.

## 4. Methods

### 4.1. The equation for the transcriptional activity regulated by the sole blocking-type repression

The transcription regulated by sole blocking-type repression (figure 2a) can be described by the following system of ordinary differential equations (ODEs) based on the mass action law:

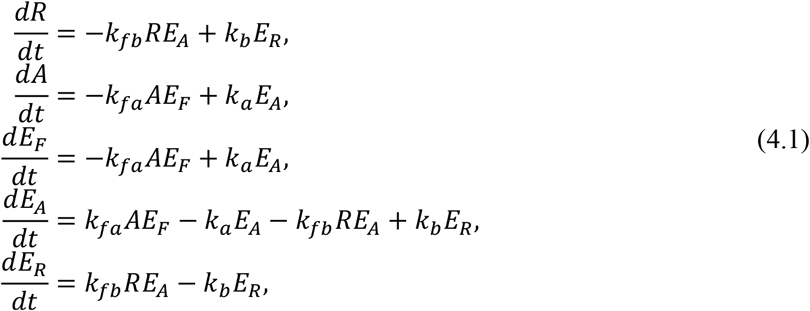

where *R, A, E_F_*, *E_A_*, and *E_R_* represent the concentration of the repressor, the activator, DNA, the activator-bound DNA, and the activator and repressor complex-bound DNA, respectively. Here, *k_fb_ (k_b_*) and *k_fa_* (*k_a_*) are the association (dissociation) rate constants between *E_A_* and *R* and between *A* and *E_F_*, respectively. Note that as 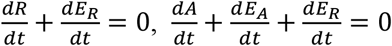, and 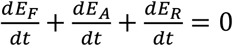, the total concentrations of the repressor (*R_T_* ≡ *R* + *E_R_*), the activator (*A_T_* ≡ *A* + *E_A_* + *E_R_*), and DNA (*E_T_* ≡ *E_F_* + *E_A_* + *E_R_*) are conserved. The steady states of the system satisfy the following equations,

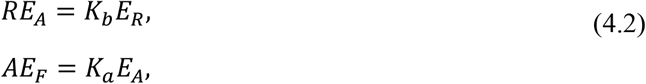

where *K_b_* = *k_b_/k_fb_* and *K_a_* = *k_a_/k_fa_*. This yields 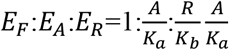, and thus the steady state for *E_A_/E_T_*:

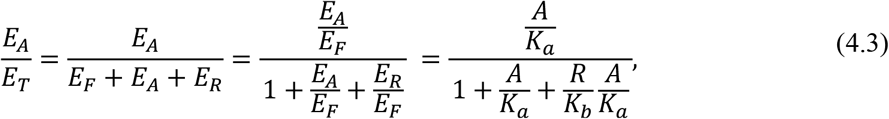

where *A* and *R* in equation (4.3) are the steady states of the free activator and the free repressor, respectively.

Equation (4.3) can be simplified if the total concentration of DNA (*E_T_*) is much lower than the concentrations of the activator (*A_T_*) and repressor (*R_T_*) and thus *A_T_* = *A* + *E_A_* + *E_R_* ≈ *A* and *R_T_* = *R* + *E_R_* ≈ *R*. That is, by replacing *A* and *R* in equation (4.3) with conserved *A_T_* and *R_T_*, respectively, we get the following approximation for equation (4.3):

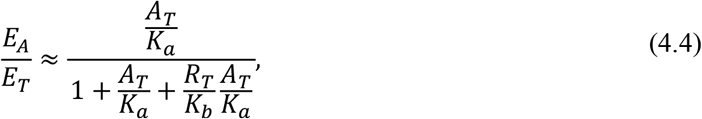

which is accurate as long as *E_T_/A_T_* is small (figure S4a). This assumption is likely to hold in the mammalian circadian clock as the number of BMAL1:CLOCK in the mammalian cells is about 10^4^~10^5^ [37]. On the other hand, it might not be acceptable in *E. coli* or *S. cerevisiae* cells, which contain much lower numbers of transcription factors (10^1^~10^2^) [37].

### 4.2. The equation for the transcriptional activity regulated by the blocking- and sequestration-type repressions

The transcription regulated by both blocking and sequestration (figure 2d) can be described by the following ODEs:

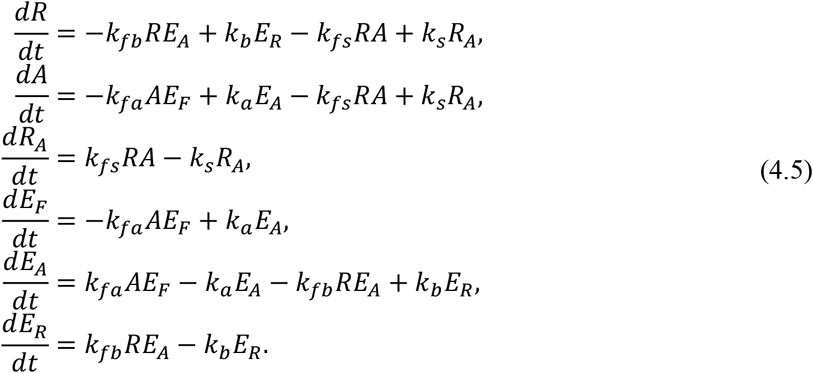

The reversible binding between *R* and *A* to form the complex (*R_A_*) with the association rate constant *k_fs_* and the dissociation rate constant *k_s_* are added to equation (4.1). Thus, the conservation laws for the activator and the repressor are changed to *A_T_* = *A* + *R_A_* + *E_A_* + *E_R_* and *R_T_* = *R* + *R_A_* + *E_R_*, respectively. Because the steady states of equation (4.5) also satisfy equation (4.2), the steady state of *E_A_/E_T_* in this system also satisfies equation (4.3). However, even if *E_T_* is much lower than *A_T_* and *R_T_*, equation (4.3) cannot be simplified by replacing *A* and *R* with *A_T_* and *R_T_* because *A_T_* = *A* + *R_A_* + *E_A_* + *E_R_* ≈ *A* + *R_A_* and *R_T_* = *R* + *R_A_* + *E_R_* ≈ *R* + *R_A_*. Thus, we also need to use another steady state equation, *AR* = *K_s_R_A_*, where *K_s_* = *k_s_/k_fs_*, to derive the steady state of *R_A_* in terms of *A_T_* and *R_T_*. Specifically, by replacing *A* and *R* with *A_T_* – *R_A_* and *R_T_* – *R_A_*, respectively, in *AR* = *K_s_R_A_*, we get 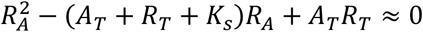, yielding the approximate steady state for *R_A_*:

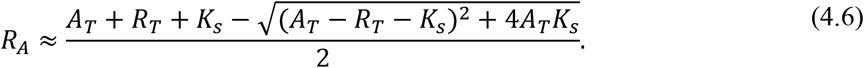

Then by substituting equation (4.6) to *A* ≈ *A_T_* – *R_A_* and *R* ≈ *R_T_* – *R_A_*, we can get the following approximate steady state for the free activator and repressor:

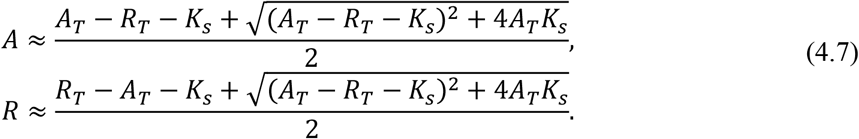

By substituting equation (4.7) to equation (4.3), the approximate *E_A_/E_T_* can be derived (equation (2.4)), which is accurate as long as *E_T_/A_T_* is small (figure S4b).

### 4.3. The equation for the transcriptional activity regulated by all the blocking-, sequestration-, and displacement-type repressions

The transcription regulated by all blocking, sequestration, and displacement (figure 2g) can be described by the following ODEs:

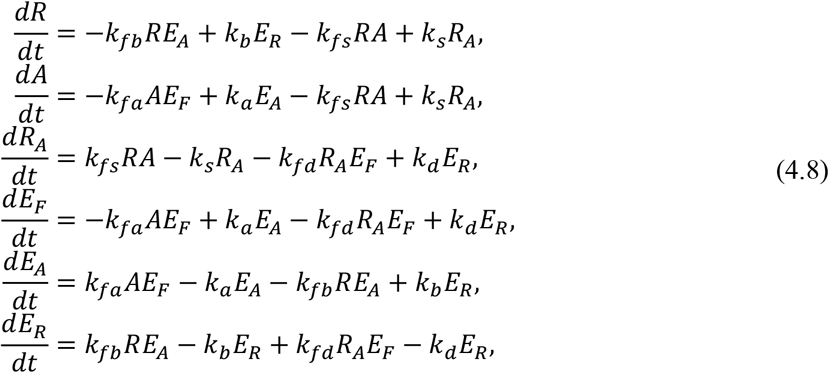

which have the same conservation laws as equation (4.5). Because the reversible binding between *R_A_* and *E_F_* to form *E_R_* with the association rate constant *k_fd_* and the dissociation rate constant *k_d_* are added to equation (4.5), the steady states are changed to the following equations:

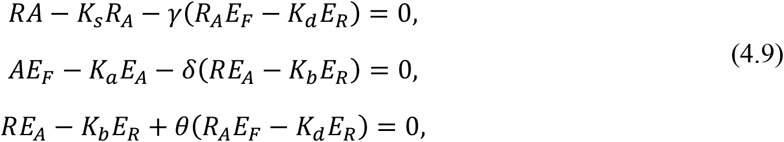

where *K_d_* = *k_d_/k_fd_*, *γ* = *k_fd_/k_fs_, δ* = *k_fb_/k_fa_*, and *θ* = *k_fd_/k_fb_*. If *E_T_* is much lower than *A_T_* and *R_T_*, and thus *A_T_* = *A* + *R_A_* + *E_A_* + *E_R_* ≈ *A* + *R_A_* and *R_T_* = *R* + *R_A_* + *E_R_* ≈ *R* + *R_A_*, the last two equations of (4.9) can be simplified as follows:

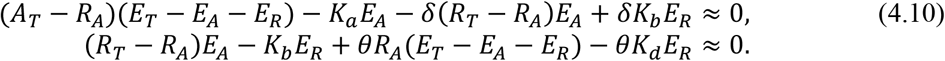

Their solution yields the steady state approximation for *E_A_/E_T_* as

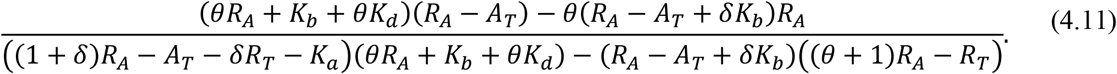

Furthermore, by using the approximation *A_T_* ≈ *A* + *R_A_* and *R_T_* ≈ *R* + *R_A_*, we can simplify the first equation of equation (4.9) as follows:

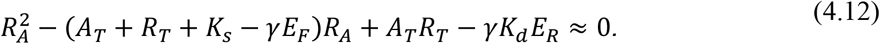

Because *E_T_* is much lower than *A_T_* and *R_T_*, equation (4.12) can be further simplified to 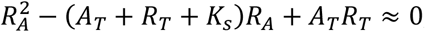, leading to the approximation for the steady state of *R_A_* described in equation (4.6). Then, by substituting equation (4.6) to equation (4.11), the approximated *E_A_/E_T_* in terms of conserved *A_T_* and *R_T_* can be derived. The approximation of *E_A_/E_T_* can be further simplified as follows if we assume *δ* = 1 and *θ* = 1 (i.e., the binding rates are the same):

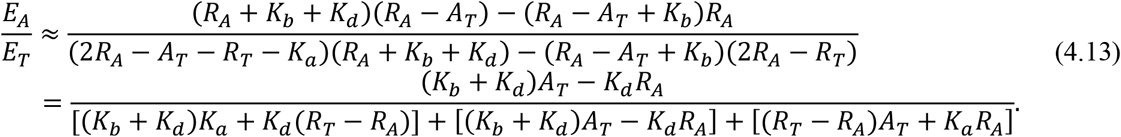

Each term of equation (4.13) can be transformed by using *A* ≈ *A_T_* – *R_A_*, *R* ≈ *R_T_* – *R_A_*, and *AR* ≈ *K_s_R_A_* as follows:

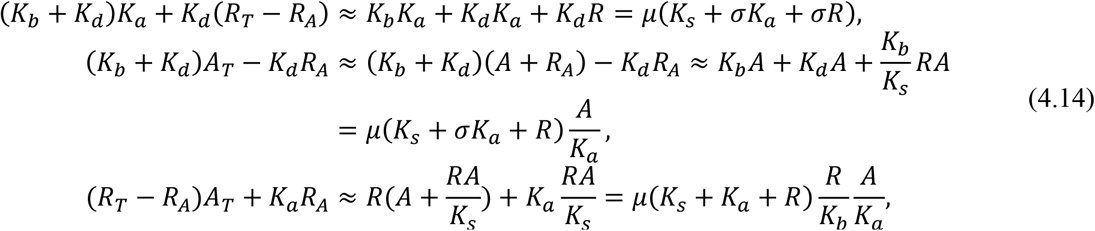

where *σ* = *K_s_K_d_/K_b_K_a_* and *μ* = *K_b_K_a_/K_s_*. By substituting equation (4.14) to equation (4.13), we can derive equation (2.5) as follows:

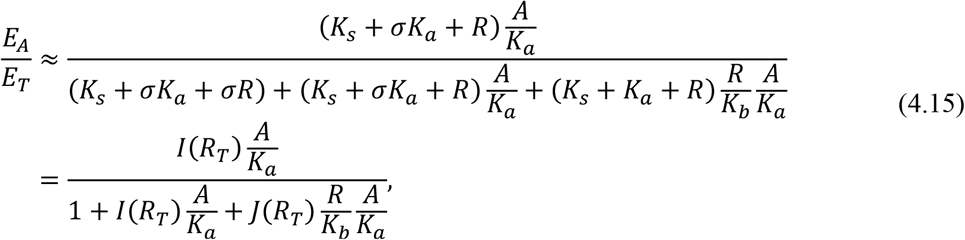

where 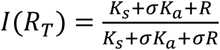 and 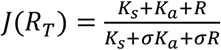. This approximation is accurate as long as *E_T_/A_T_* is small (figure S4c). Importantly, it accurately captures the cases when *δ* and *θ* are not one if the displacement effectively occurs (*K_d_* > *K_a_*; figure S5).

