## Supplementary material for "Combined multiple transcriptional repression mechanisms generate ultrasensitivity and oscillations": SI Appendix

### 1. The transcriptional NFL model

The transcriptional NFL model incorporating blocking-, sequestration-, and displacement-type repressions (figure 3a) is described by the following system of ODEs based on mass-action kinetics:

$$\begin{aligned}
\frac{dM}{dt} &= \alpha_1 \frac{E_A}{E_T} - \beta M, \\
\frac{dR_c}{dt} &= \alpha_2 M - \beta R_c, \\
\frac{dR}{dt} &= \alpha_3 R_c - \beta R - k_{fb} R E_A + k_b E_R - k_{fs} R A + k_s R_A, \\
\frac{dA}{dt} &= -k_{fa} A E_F + k_a E_A - k_{fs} R A + k_s R_A + \beta R_A, \\
\frac{dR_A}{dt} &= k_{fs} R A - k_s R_A - \beta R_A - k_{fd} R_A E_F + k_d E_R, \\
\frac{dE_F}{dt} &= -k_{fa} A E_F + k_a E_A - k_{fd} R_A E_F + k_d E_R, \\
\frac{dE_A}{dt} &= k_{fa} A E_F - k_a E_A - k_{fb} R E_A + k_b E_R + \beta E_R, \\
\frac{dE_R}{dt} &= k_{fb} R E_A - k_b E_R + k_{fd} R_A E_F - k_d E_R - \beta E_R.
\end{aligned} \tag{S1}$$

Here, $M$, $R_c$, $R$, $A$, $R_A$, $E_F$, $E_A$, and $E_R$ represent the concentration of the repressor mRNA, the repressor in the cytoplasm, the repressor in the nucleus, the activator, the repressor and activator complex, the free DNA, the activator-bound DNA, and the activator and repressor complex-bound DNA, respectively. $M$ is transcribed with the rate of $\alpha_1$ and then translated to the repressor protein $(R_c)$ in the cytoplasm with the rate of $\alpha_2$. The repressor $(R)$ translocated into the nucleus with the rate of $\alpha_3$ inhibits the activator $(A)$ with blocking, sequestration, and displacement. We assume that the degradation rates of the mRNA and the repressor are the same as $\beta$ to increase the chance of oscillations [1-3].

As the total concentrations of the activator $(A_T \equiv A + R_A + E_A + E_R)$ and DNA $(E_T \equiv E_F + E_A + E_R)$ are conserved (i.e., $\frac{dA}{dt} + \frac{dR_A}{dt} + \frac{dE_A}{dt} + \frac{dE_R}{dt} = 0$ and $\frac{dE_F}{dt} + \frac{dE_A}{dt} + \frac{dE_R}{dt} = 0$), we can simplify the model as follows:

$$\begin{aligned}
\frac{dM}{dt} &= \alpha_1 \frac{E_A}{E_T} - \beta M, \\
\frac{dR_c}{dt} &= \alpha_2 M - \beta R_c, \\
\frac{dR}{dt} &= \alpha_3 R_c - \beta R - k_{fb} R E_A + k_b E_R - k_{fs} R (A_T - R_A - E_A - E_R) + k_s R_A, \\
\frac{dR_A}{dt} &= k_{fs} R (A_T - R_A - E_A - E_R) - k_s R_A - \beta R_A - k_{fd} R_A (E_T - E_A - E_R) + k_d E_R, \\
\frac{dE_A}{dt} &= k_{fa} (A_T - R_A - E_A - E_R)(E_T - E_A - E_R) - k_a E_A - k_{fb} R E_A + k_b E_R + \beta E_R, \\
\frac{dE_R}{dt} &= k_{fb} R E_A - k_b E_R + k_{fd} R_A (E_T - E_A - E_R) - k_d E_R - \beta E_R.
\end{aligned} \tag{S2}$$

By non-dimensionalizing the model with the scaling of variables as $M = \rho \frac{\beta^2}{\alpha_2 \alpha_3} M^*$, $R_c = \rho \frac{\beta}{\alpha_3} R_c^*$, and $X = \rho X^*$ $(X = R, A, R_A, A_T, E_F, E_A, E_R, E_T)$, with the scaling of binding rates as $k_{fi} = \frac{\beta}{\rho} k_{fi}^*$ $(i = b, a, s, d)$, with the scaling of unbinding rates as $k_i = \beta k_i^*$ $(i = b, a, s, d)$, and with the scaling of time as $t = \frac{t^*}{\beta}$, where $\rho = \frac{\alpha_1 \alpha_2 \alpha_3}{\beta^3}$, we can further simplify the model as follows:

$$
\begin{aligned}
\frac{dM^*}{dt^*} &= \frac{E_A^*}{E_T^*} - M^*, \\
\frac{dR_c^*}{dt^*} &= M^* - R_c^*, \\
\frac{dR^*}{dt^*} &= R_c^* - R^* - k_{fs}^* R^* (A_T^* - R_A^* - E_A^* - E_R^*) + k_s^* R_A^* - k_{fb}^* R^* E_A^* + k_b^* E_R^* \\
\frac{dR_A^*}{dt^*} &= k_{fs}^* R^* (A_T^* - R_A^* - E_A^* - E_R^*) - k_s^* R_A^* - R_A^* - k_{fd}^* R_A^* (E_T^* - E_A^* - E_R^*) + k_d^* E_R^*, \\
\frac{dE_A^*}{dt^*} &= k_{fa}^* (A_T^* - R_A^* - E_A^* - E_R^*)(E_T^* - E_A^* - E_R^*) - k_a^* E_A^* - k_{fb}^* R^* E_A^* + k_b^* E_R^* + E_R^*, \\
\frac{dE_R^*}{dt^*} &= k_{fd}^* R_A^* (E_T^* - E_A^* - E_R^*) - k_d^* E_R^* + k_{fb}^* R^* E_A^* - k_b^* E_R^* - E_R^*,
\end{aligned}
\tag{S3}
$$

where $A_T^* = A^* + R_A^* + E_A^* + E_R^*$, and $E_T^* = E_F^* + E_A^* + E_R^*$. For simplicity, we omit * in the notation of the variables and parameters (e.g., we use $M$ instead of $M^*$) from here.

To further simplify the model, we will utilize the fact that the reversible bindings are typically much faster than the other reactions (i.e., $k_b, k_a, k_s, k_d, k_{fb}, k_{fa}, k_{fs}, k_{fd} \gg 1$). In this case, $M$ and $R_c$ are slow variables affected only by slow reactions, not fast reversible binding reactions. On the other hand, $R$ is affected by both slow and fast reactions. Thus, we replace $R$ with the total repressor concentration in the nucleus ($R_T \equiv R + R_A + E_R$), which is affected only by the slow reactions because $\frac{dR_T}{dt} = \frac{dR}{dt} + \frac{dR_A}{dt} + \frac{dE_R}{dt} = R_c - R_T$ (see [4] for details). By substituting $R_T - R_A - E_R$ for $R$ in equation S3, we can get the model with the complete time scale separation among variables (i.e., $M$, $R_c$, and $R_T$ are slow variables and the others are fast variables) as follows:

$$
\begin{aligned}
\frac{dM}{dt} &= \frac{E_A}{E_T} - M, \\
\frac{dP_c}{dt} &= M - R_c, \\
\frac{dR_T}{dt} &= R_c - R_T, \\
\frac{dR_A}{dt} &= k_{fs} R (A_T - R_A - E_A - E_R) - (k_s R_A + R_A) - k_{fd} R_A (E_T - E_A - E_R) + k_d E_R, \\
\frac{dE_A}{dt} &= k_{fa} (A_T - R_A - E_A - E_R)(E_T - E_A - E_R) - k_a E_A - k_{fb} R E_A + (k_b E_R + E_R), \\
\frac{dE_R}{dt} &= k_{fd} R_A (E_T - E_A - E_R) - k_d E_R + k_{fb} R E_A - (k_b E_R + E_R).
\end{aligned}
\tag{S4}
$$

Using the fact that the fast variables $R_A$, $E_A$, and $E_R$ reach their quasi-steady-state rapidly compared to the slow variables $M$, $R_c$, and $R_T$, we can eliminate those variables by replacing them with their quasi-steady-state approximations (QSSAs). The QSSAs can be obtained by solving $\frac{dR_A}{dt} = 0$, $\frac{dE_A}{dt} =$

$0$, and $\frac{dE_R}{dt} = 0$. These equations are nearly identical with the steady state equations of equation (4.8) because $k_s R_A + R_A \approx k_s R_A$ in $\frac{dR_A}{dt}$ and $k_b E_R + E_R \approx k_b E_R$ in $\frac{dE_A}{dt}$ and $\frac{dE_R}{dt}$. Thus, we can use equation (2.5) as the QSSA for $E_A/E_T$. This allows us to get the simplified transcriptional NFL model as follows:

$$\frac{dM}{dt} = \frac{E_A(\tilde{R}_T)}{E_T} - M,$$
$$\frac{dR_c}{dt} = M - R_c,$$
$$\frac{dR_T}{dt} = R_c - R_T,$$

where $E_A(\tilde{R}_T)/E_T$ is equation (2.5). Similarly, using equations (2.2) and (2.4) for $E_A(\tilde{R}_T)/E_T$, we can obtain the transcriptional NFL model with the sole blocking and the blocking and sequestration, respectively.

### 2. The equations for the transcriptional activity regulated by the combinations of sequestration and displacement, and blocking and displacement

#### 2.1. The equation for the transcriptional activity regulated by the sequestration- and displacement-type repressions

The transcription regulated by both sequestration and displacement (figure S3a) can be described by the following ODEs:

$$\begin{aligned}
\frac{dR}{dt} &= -k_{fs}RA + k_sR_A,\\
\frac{dA}{dt} &= -k_{fs}RA + k_sR_A - k_{fa}AE_F + k_aE_A,\\
\frac{dR_A}{dt} &= k_{fs}RA - k_sR_A - k_{fd}R_AE_F + k_dE_R,\\
\frac{dE_F}{dt} &= -k_{fa}AE_F + k_aE_A - k_{fd}R_AE_F + k_dE_R,\\
\frac{dE_A}{dt} &= k_{fa}AE_F - k_aE_A,\\
\frac{dE_R}{dt} &= k_{fd}R_AE_F - k_dE_R,
\end{aligned} \tag{S5}$$

where $R$, $A$, $E_F$, $E_A$, and $E_R$ represent the concentrations of the repressor, the activator, the free DNA, the activator-bound DNA, and the activator and repressor complex-bound DNA, respectively. Here, $k_{fs}$ ($k_s$), $k_{fa}$ ($k_a$), and $k_{fd}$ ($k_d$) are the association (dissociation) rate constants between $A$ and $R$, $A$ and $E_F$, and $R_A$ and $E_F$, respectively. Note that as $\frac{dR}{dt}+\frac{dR_A}{dt}+\frac{dE_R}{dt}=0$, $\frac{dA}{dt}+\frac{dR_A}{dt}+\frac{dE_A}{dt}+\frac{dE_R}{dt}=0$, and $\frac{dE_F}{dt}+\frac{dE_A}{dt}+\frac{dE_R}{dt}=0$, the total concentrations of the repressor ($R_T \equiv R + R_A + E_R$), the activator ($A_T \equiv A + R_A + E_A + E_R$), and DNA ($E_T \equiv E_F + E_A + E_R$) are conserved. The steady states of the system satisfy the following equations:

$$\begin{aligned}
AE_F &= K_aE_A,\\
R_AE_F &= K_dE_R,
\end{aligned} \tag{S6}$$

where $K_a = k_a/k_{fa}$ and $K_d = k_d/k_{fd}$. This yields $E_F{:}E_A{:}E_R = 1{:}\frac{A}{K_a}{:}\frac{R_A}{K_d}$, and thus the steady state for $E_A/E_T$:

$$\frac{E_A}{E_T} = \frac{E_A}{E_F+E_A+E_R} = \frac{\frac{E_A}{E_F}}{1+\frac{E_A}{E_F}+\frac{E_R}{E_F}} = \frac{\frac{A}{K_a}}{1+\frac{A}{K_a}+\frac{R_A}{K_d}}, \tag{S7}$$

where $A$ and $R$ in equation (S7) are the steady states of the free activator and the free repressor, respectively.

To derive the steady states of $A$ and $R$ in terms of $A_T$ and $R_T$, we use another steady state equation $AR = K_sR_A$, where $K_s = k_s/k_{fs}$. Because $E_T$ is much lower than $A_T$ and $R_T$, and thus

$A_T = A + R_A + E_A + E_R \approx A + R_A$ and $R_T = R + R_A + E_R \approx R + R_A$, we replace $A$ and $R$ in $AR = K_s R_A$ with $A_T - R_A$ and $R_T - R_A$, respectively. Then, we get $R_A^2 - (A_T + R_T + K_s)R_A + A_T R_T \approx 0$, yielding the approximate steady state for $R_A$:

$$R_A \approx \frac{A_T + R_T + K_s - \sqrt{(A_T - R_T - K_s)^2 + 4A_T K_s}}{2}. \tag{S8}$$

Then by substituting equation (S8) to $A \approx A_T - R_A$, we can get the following approximate steady state for the free activator:

$$A \approx \frac{A_T - R_T - K_s + \sqrt{(A_T - R_T - K_s)^2 + 4A_T K_s}}{2}. \tag{S9}$$

By substituting equations (S8) and (S9) to equation (S7), the approximate $E_A/E_T$ can be derived, which is accurate as long as $E_T/A_T$ is small. Note that equation (S7) is equivalent to equation (4.15) if $\sigma = \frac{K_s K_d}{K_b K_a} = 1$ because $AR = K_s R_A$ (i.e., $\frac{R_A}{K_d} = \frac{R}{K_s}\frac{A}{K_d} = \frac{R}{K_b}\frac{A}{K_a}$). That is, the transcriptional activity regulated by the three repressions is equivalent to that by the sequestration and displacement under the detailed balance condition.

##### 2.2 The equation for the transcriptional activity regulated by the blocking- and displacement-type repressions

The transcription regulated by both blocking and displacement (figure S3b) can be described by the following ODEs:

$$\begin{aligned}
\frac{dR}{dt} &= -k_{fb} R E_A + k_b E_R, \\
\frac{dA}{dt} &= -k_{fa} A E_F + k_a E_A, \\
\frac{dR_A}{dt} &= -k_{fd} R_A E_F + k_d E_R, \\
\frac{dE_F}{dt} &= -k_{fa} A E_F + k_a E_A - k_{fd} R_A E_F + k_d E_R, \\
\frac{dE_A}{dt} &= k_{fa} A E_F - k_a E_A - k_{fb} R E_A + k_b E_R, \\
\frac{dE_R}{dt} &= k_{fb} R E_A - k_b E_R + k_{fd} R_A E_F - k_d E_R.
\end{aligned} \tag{S10}$$

The reversible binding between $R_A$ and $E_F$ to form the ternary complex ($E_R$) with the association rate constant $k_{fd}$ and the dissociation rate constant $k_d$ are added to equation (4.1). Note that as $\frac{dR}{dt} + \frac{dR_A}{dt} + \frac{dE_R}{dt} = 0$, $\frac{dA}{dt} + \frac{dR_A}{dt} + \frac{dE_A}{dt} + \frac{dE_R}{dt} = 0$, and $\frac{dE_F}{dt} + \frac{dE_A}{dt} + \frac{dE_R}{dt} = 0$, the total concentrations of the repressor ($R_T \equiv R + R_A + E_R$), the activator ($A_T \equiv A + R_A + E_A + E_R$), and DNA ($E_T \equiv E_F + E_A + E_R$) are conserved. The steady states of the system satisfy the following equations:

$$\begin{aligned}
R E_A &= K_b E_R, \\
A E_F &= K_a E_A, \\
R_A E_F &= K_d E_R,
\end{aligned} \tag{S11}$$

where $K_b = k_b/k_{fb}$, $K_a = k_a/k_{fa}$, and $K_d = k_d/k_{fd}$. This yields $E_F : E_A : E_R = 1 : \frac{A}{K_a} : \frac{R_A}{K_d} = 1 : \frac{A}{K_a} : \frac{R}{K_b}\frac{A}{K_a}$, and thus the steady state for $E_A/E_T$:

$$\frac{E_A}{E_T} = \frac{E_A}{E_F + E_A + E_R} = \frac{\frac{E_A}{E_F}}{1 + \frac{E_A}{E_F} + \frac{E_R}{E_F}} = \frac{\frac{A}{K_a}}{1 + \frac{A}{K_a} + \frac{R_A}{K_d}} = \frac{\frac{A}{K_a}}{1 + \frac{A}{K_a} + \frac{R}{K_b}\frac{A}{K_a}}, \tag{S12}$$

where $A$ and $R$ in equation (S12) are the steady states of the free activator and the free repressor, respectively.

To derive the steady states of $A$ and $R$ in terms of $A_T$ and $R_T$, we use the relation $\frac{E_R}{E_F} = \frac{R_A}{K_d} = \frac{R}{K_b}\frac{A}{K_a}$, that is $AR = \frac{K_b K_a}{K_d} R_A$. Because $E_T$ is much lower than $A_T$ and $R_T$, and thus $A_T = A + R_A + E_A + E_R \approx A + R_A$ and $R_T = R + R_A + E_R \approx R + R_A$, we replace $A$ and $R$ in $AR = \frac{K_b K_a}{K_d} R_A$ with $A_T - R_A$ and $R_T - R_A$, respectively. Then we can obtain the approximate steady states of $A$ and $R$:

$$A \approx \frac{A_T - R_T - \frac{K_b K_a}{K_d} + \sqrt{\left(A_T - R_T - \frac{K_b K_a}{K_d}\right)^2 + 4A_T \frac{K_b K_a}{K_d}}}{2}, \qquad R \approx \frac{R_T - A_T - \frac{K_b K_a}{K_d} + \sqrt{\left(A_T - R_T - \frac{K_b K_a}{K_d}\right)^2 + 4A_T \frac{K_b K_a}{K_d}}}{2}. \tag{S13}$$

By substituting equation (S13) to equation (S12), the approximate $E_A/E_T$ can be derived, which is accurate as long as $E_T/A_T$ is small. Note that equations (S12) and (S13) are equivalent to equations (4.15) and (4.7), respectively, if $\sigma = \frac{K_s K_d}{K_b K_a} = 1$. That is, the transcriptional activity regulated by all three repressions is equivalent to that by blocking and displacement under the detailed balance condition.

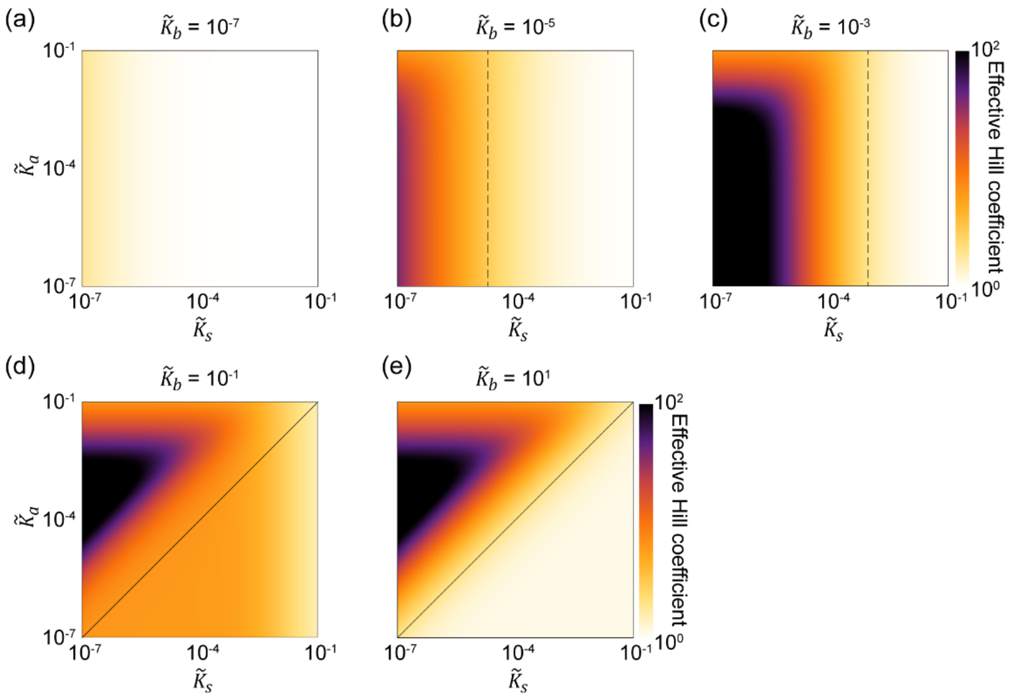


**Figure S1. The combination of sequestration and blocking can generate ultrasensitivity when they act synergistically. (a)** When the blocking is too strong compared to the sequestration ($\widetilde{K}_s > \widetilde{K}_b$), it is challenging to generate ultrasensitivity, similar to the sole blocking, as shown in figure 2c. **(b-c)** As the blocking becomes weaker than the sequestration (i.e., $\widetilde{K}_s < \widetilde{K}_b$; left of the dashed lines), ultrasensitivity can be generated. **(d-e)** When the blocking becomes too weak ($\widetilde{K}_s \ll \widetilde{K}_b$), and thus sequestration dominates, ultrasensitivity is generated in a more limited region. In particular, ultrasensitivity cannot be generated if the activator binds to DNA more tightly than the repressor (i.e., $\widetilde{K}_s < \widetilde{K}_a$; below the solid lines).

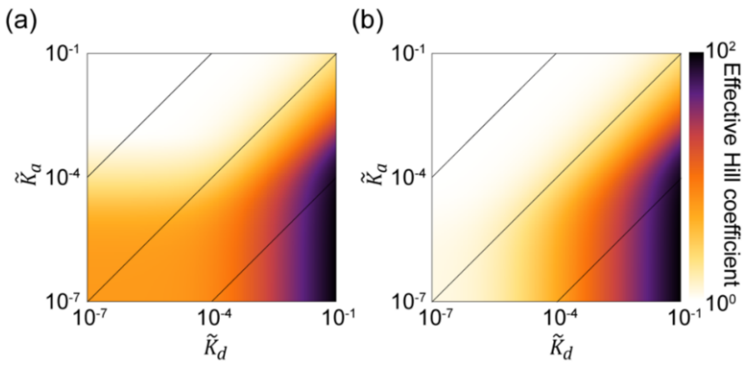


**Figure S2. Even if $\widetilde{K}_b \neq \widetilde{K}_s$, ultrasensitivity can be generated when effective displacement occurs. (a-b)** Both when the sequestration is stronger than the blocking ($\widetilde{K}_b = 10^{-4}$ and $\widetilde{K}_s = 10^{-5}$; (a)) and when the blocking is stronger than the sequestration ($\widetilde{K}_b = 10^{-6}$ and $\widetilde{K}_s = 10^{-5}$; (a)), ultrasensitivity can be generated as the effective displacement occurs (i.e., $\widetilde{K}_d \gg \widetilde{K}_a$).

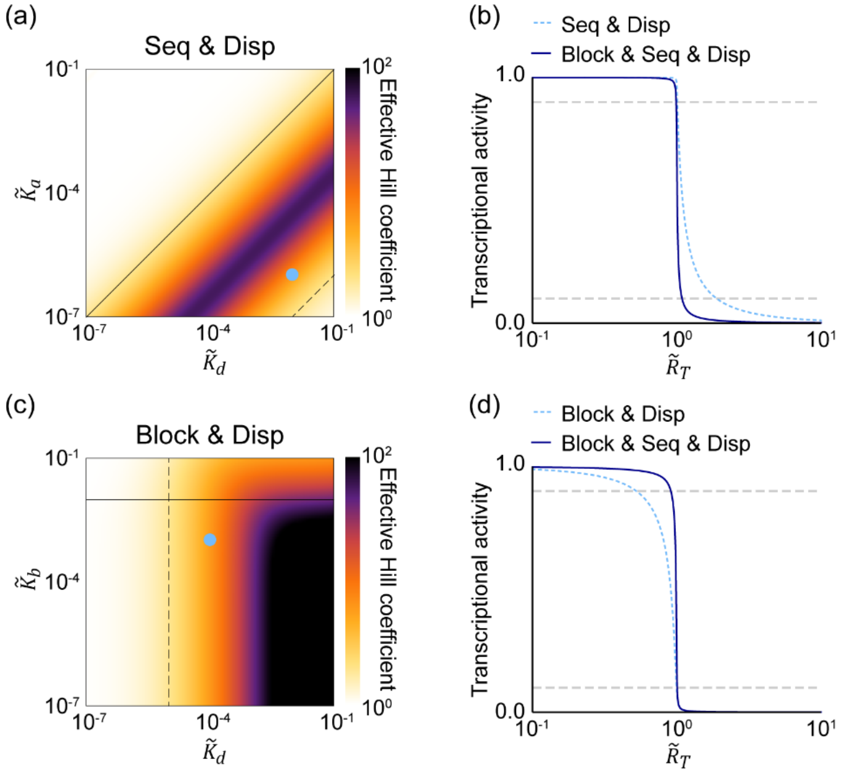


**Figure S3. When the additional repression mechanism is added to the combinations of sequestration and displacement repressions and blocking and displacement repressions, more sensitive transcription response can be generated. (a)** The heat map represents the effective Hill coefficient of the transcriptional activity regulated by the combination of the sequestration- and displacement-type repressions (equation (S7)). For the combination of sequestration and displacement repressions to generate the ultrasensitivity of the transcription response, the effective displacement must occur (i.e., $\tilde{K}_d/\tilde{K}_a > 1$; below the solid lines), but become weaker than the sequestration (i.e., $\tilde{K}_d/\tilde{K}_a \ll 1/\tilde{K}_s$; above the dashed lines). Here, $\tilde{K}_s = 10^{-5}$. **(b)** With the parameter ($\tilde{K}_d = 10^{-2}, \tilde{K}_a = 10^{-6}, \tilde{K}_s = 10^{-5}$; circle mark in (a)), the combination of the sequestration and displacement fails to generate ultrasensitivity (dotted line). When the blocking is added, the ultrasensitivity is generated (solid line). Here, $\tilde{K}_b = 10^{-2}$ was chosen to break the detailed balanced condition. **(c)** The heat map represents the effective Hill coefficient of the transcriptional activity regulated by the combination of the blocking- and displacement-type repressions (equation (S12)). For the combination of blocking and displacement repressions to generate the ultrasensitivity of the transcription response, the blocking should not be too weak (i.e., $\tilde{K}_b < 10^{-2}$; below the solid line), and the effective displacement must occur (i.e., $\tilde{K}_a < \tilde{K}_d$; right of the dashed line). Here, $\tilde{K}_a = 10^{-5}$. **(d)** With the parameter ($\tilde{K}_d = 10^{-4}, \tilde{K}_b = 10^{-3}, \tilde{K}_a = 10^{-5}$; circle mark in (c)), the combination of the blocking and displacement fails to generate ultrasensitivity (dotted line). When the sequestration is added, the ultrasensitivity is generated (solid line). Here, $\tilde{K}_s = 10^{-5}$ was chosen to break the detailed balanced condition.

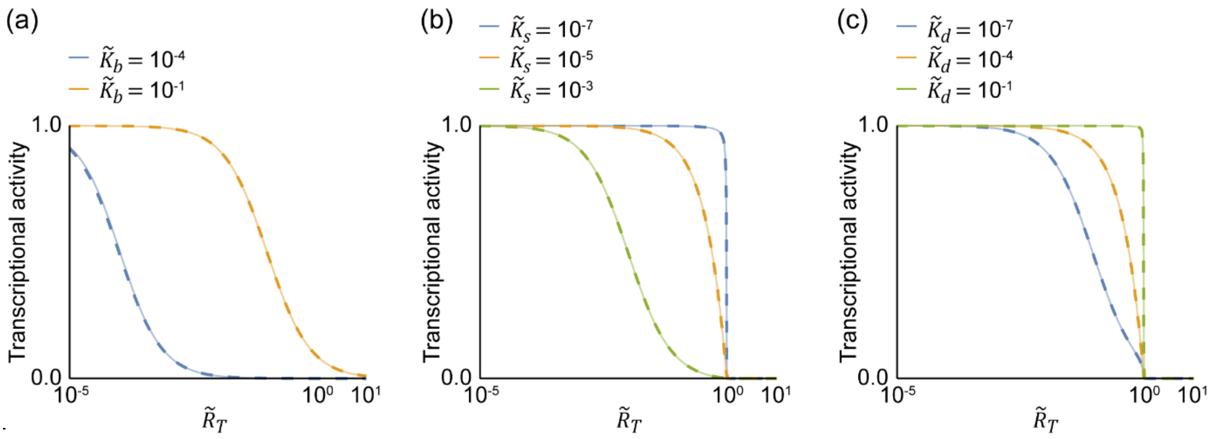


**Figure S4. When the total concentration of DNA is much lower than that of the activator, the approximated transcriptional activity is accurate. (a-c)** We can simplify the steady state equations for $E_A/E_T$ in the systems of equations (4.1), (4.5), and (4.8) by assuming that the total concentration of DNA is much lower than that of the activator (i.e., $E_T/A_T \ll 1$). Indeed, when $E_T/A_T = 10^{-5}$, the exactly solved transcriptional activity (solid lines) is accurately approximated by the simplified transcriptional activity (dashed lines) regulated by blocking (a), blocking and sequestration (b), and blocking, sequestration, and displacement (c). The dashed lines in (a), (b), and (c) came from figure 2b, e, and h, respectively.

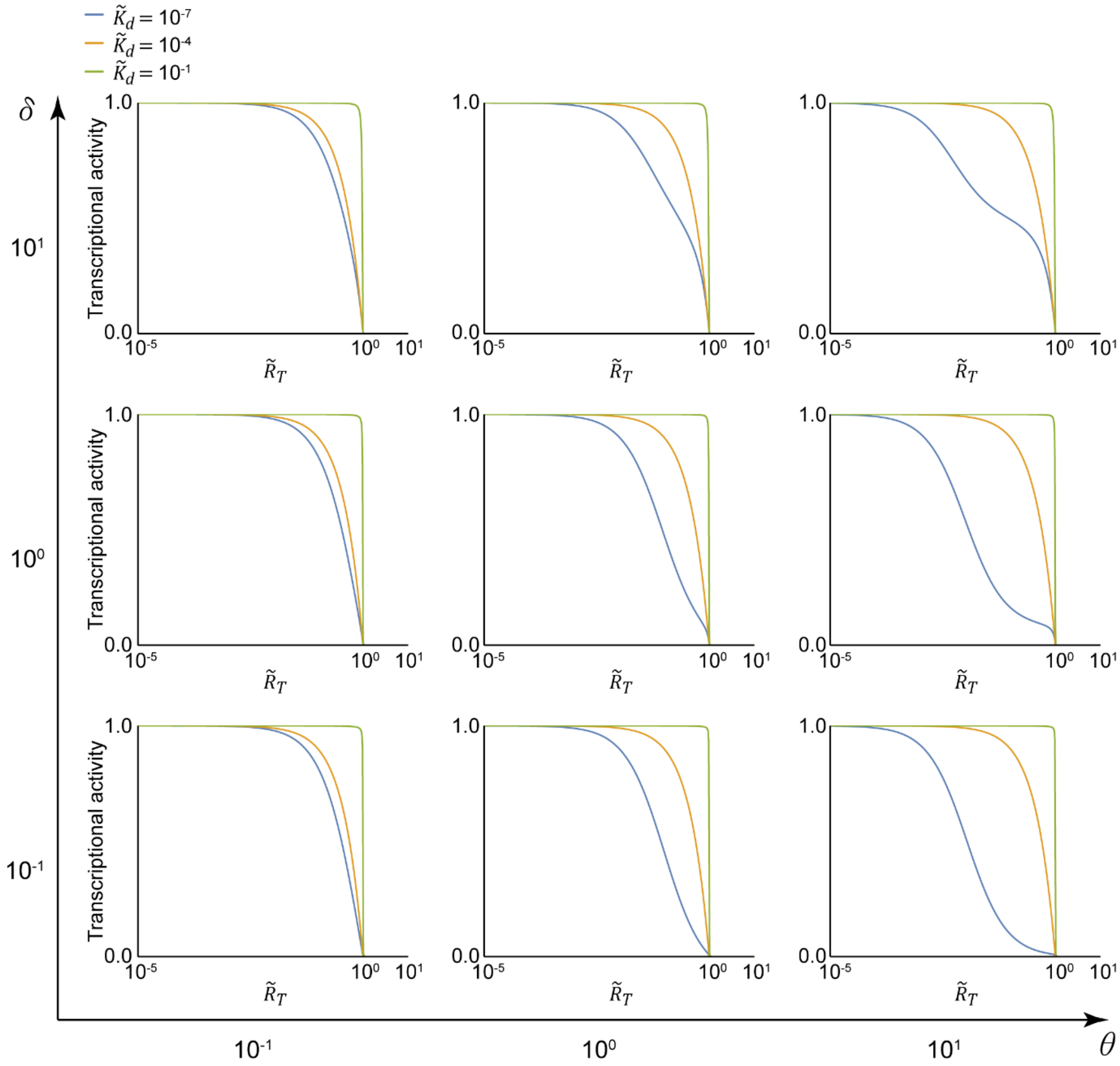


**Figure S5. Effective displacement can generate ultrasensitivity regardless of the disparity in the binding rates of the activator, the repressor, and the activator-repressor complex to DNA.** As the ratio between the binding rates of the repressor to the DNA-bound activator and the activator to DNA ($\delta = k_{fb}/k_{fa}$), and the ratio between the binding rates of the repressor-activator complex to DNA and the repressor to the DNA-bound activator ($\theta = k_{fd}/k_{fb}$) vary, the transcriptional activity changes slightly when the effective displacement does not occur (i.e., $\tilde{K}_d = 10^{-7} < \tilde{K}_a$; blue lines). On the other hand, as the displacement becomes effective (i.e., $\tilde{K}_d = 10^{-1} > \tilde{K}_a$; green lines), the transcriptional activity is nearly the same regardless of $\delta$ and $\theta$, indicating that strong displacement can generate ultrasensitivity regardless of $\delta$ and $\theta$ (green lines). Here $\tilde{K}_a = 10^{-4}$.

| Blocking, Sequestration, and Displacement | | |
|---|---|---|
| $\tilde{K}_b \ll 1,\ \tilde{K}_s \ll 1,\ \tilde{K}_d/\tilde{K}_a > 1$ (figure 2 and S2) | | |
| **Blocking and Sequestration** | **Sequestration and Displacement** | **Blocking and Displacement** |
| $\tilde{K}_b \ll 1,\ \tilde{K}_s \ll 1,\ \tilde{K}_d/\tilde{K}_a > 1$ with $\tilde{K}_d = \tilde{K}_b\tilde{K}_a/\tilde{K}_s$ $\Leftrightarrow \tilde{K}_s < \tilde{K}_b \ll 1$ (figure 2f and S1) | $\tilde{K}_b \ll 1,\ \tilde{K}_s \ll 1,\ \tilde{K}_d/\tilde{K}_a > 1$ with $\tilde{K}_b = \tilde{K}_s\tilde{K}_d/\tilde{K}_a$ $\Leftrightarrow 1 < \tilde{K}_d/\tilde{K}_a \ll 1/\tilde{K}_s$ (figure S3a) | $\tilde{K}_b \ll 1,\ \tilde{K}_s \ll 1,\ \tilde{K}_d/\tilde{K}_a > 1$ with $\tilde{K}_s = \tilde{K}_b\tilde{K}_a/\tilde{K}_d$ $\Leftrightarrow \tilde{K}_b \ll 1,\ \tilde{K}_d/\tilde{K}_a > 1$ (figure S3b) |

**Table S1. Conditions for ultrasensitivity generation by all combinations of blocking, sequestration, and displacement.** When all blocking, sequestration, and displacement repressions effectively occur (i.e., $\tilde{K}_b \ll 1,\ \tilde{K}_s \ll 1,\ \tilde{K}_d/\tilde{K}_a > 1$), the ultrasensitive transcription response is generated. Under the detailed balance condition (i.e., $\frac{\tilde{K}_s\tilde{K}_d}{\tilde{K}_b\tilde{K}_a} = 1$), the transcription becomes equivalent to the transcription regulated by any combination of two among blocking, sequestration, and displacement repressions (see Section 2). Thus, the transcriptions regulated by two repressions generate the ultrasensitivity under the condition that the dissociation constants of the other repression in $\tilde{K}_b \ll 1,\ \tilde{K}_s \ll 1,\ \tilde{K}_d/\tilde{K}_a > 1$ is replaced by using $\frac{\tilde{K}_s\tilde{K}_d}{\tilde{K}_b\tilde{K}_a} = 1$.
